# Rescue of *Escherichia coli* auxotrophy by *de novo* small proteins

**DOI:** 10.1101/2022.04.13.488163

**Authors:** Arianne M. Babina, Serhiy Surkov, Weihua Ye, Jon Jerlström-Hultqvist, Mårten Larsson, Erik Holmqvist, Per Jemth, Dan I. Andersson, Michael Knopp

## Abstract

Increasing numbers of small proteins with diverse physiological roles are being identified and characterized in both prokaryotic and eukaryotic systems, but the origins and evolution of these proteins remain unclear. Recent genomic sequence analyses in several organisms suggest that new functions encoded by small open reading frames (sORFs) may emerge *de novo* from noncoding sequences. However, experimental data demonstrating if and how randomly generated sORFs can confer beneficial effects to cells are limited. Here we show that by up-regulating *hisB* expression, *de novo* small proteins (≤ 50 amino acids in length) selected from random sequence libraries can rescue *Escherichia coli* cells that lack the conditionally essential SerB enzyme. The recovered small proteins are hydrophobic and confer their rescue effect by binding to the 5’ end regulatory region of the *his* operon mRNA, suggesting that protein binding promotes structural rearrangements of the RNA that allow increased *hisB* expression. This study adds RNA regulatory elements as another interacting partner for *de novo* proteins isolated from random sequence libraries, and provides further experimental evidence that small proteins with selective benefits can originate from the expression of nonfunctional sequences.

## INTRODUCTION

Once overlooked, the study of small proteins is a rapidly growing field. Typically defined as polypeptides consisting of 50 or fewer amino acids, small proteins originate from the translation of distinct small open reading frames (sORFs), rather than from the cleavage of larger precursor proteins or synthesis via ribosome-independent mechanisms (Hemm, Weaver, and Storz 2020; Storz, Wolf, and Ramamurthi 2014). Recent advancements in genome and transcriptome sequencing, ribosome-profiling techniques, proteomics, and bioinformatic analyses have led to the discovery of numerous previously unannotated small proteins in all domains of life and efforts to elucidate the targets and functions of these proteins (Andrews and Rothnagel 2014; D’Lima et al. 2017; Hemm et al. 2008; Steinberg and Koch 2021; Su et al. 2013; Weaver et al. 2019; Weidenbach et al. 2021; Yuan, D’Lima, and Slavoff 2018). The detection and characterization of small proteins in bacteria have been a particularly prolific research area over the past decade, and bacterial small proteins have been implicated in many fundamental physiological processes, including cell division, sporulation, lysis, transport, stress responses, virulence, antibiotic resistance, and cell-to-cell communication (for reviews, see: (Duval and Cossart 2017; Garai and Blanc-Potard 2020; Hemm et al. 2020; Storz et al. 2014)). Nevertheless, despite the progress made in the identification and validation of an increasing number of small proteins with versatile cellular functions, much remains unknown about the origins, evolution, and phylogenetic distribution of these small genes and their encoded proteins.

In addition to the current underannotation of sORFs within genomic databases, the short length of the coding sequences and subsequent lack of conserved protein domains render it challenging to identify small protein orthologs and establish evolutionary relationships between small proteins across different organisms. For select bacterial small proteins encoded within operons containing larger, more conserved proteins, conservation of gene synteny and/or operon content has aided in the identification of orthologs in other bacteria (Horler and Vanderpool 2009; Storz et al. 2014), but only a handful of bacterial small proteins have been found to traverse multiple phylogenetic classes. Instead, most appear to be poorly conserved and are limited to a single species or a few closely-related bacteria (Alix and Blanc-Potard 2009; Storz et al. 2014). This lack of conservation raises the question: where did these small protein-coding genes come from? Are they *bona fide* new genes that emerged independently or the remnants of genes that once encoded larger proteins?

A plausible mechanism for the *de novo* emergence of small protein-coding genes is the proto-gene model, wherein the transcription of noncoding DNA and subsequent ribosome association lead to the synthesis of novel proteins. Large-scale expression studies demonstrate pervasive transcription of non-genic stretches within characterized genomes (Dinger et al. 2008) and ribosome-profiling data indicate that many noncoding RNAs are engaged by the ribosome (Carvunis et al. 2012; Wilson and Masel 2011), supporting the notion that the expression of randomly occurring sORFs from non-genic sequences can serve as a pool for the *de novo* selection of beneficial functions (Baek et al. 2017; Hemm et al. 2008; Samayoa, Yildiz, and Karplus 2011). Furthermore, several reports show that genes recently emerged from noncoding DNA are often short in length, poorly conserved, composed primarily of hydrophobic amino acids, and tend to form alpha-helical domains – all of which are hallmark characteristics of most small proteins described to date (Carvunis et al. 2012; Storz et al. 2014).

Recent experimental work from our group has demonstrated that *de novo* small proteins with beneficial functions can be selected *in vivo* from completely random DNA sequence libraries. Due to their small size, the functions of the proteins recovered from these studies, as well as those of most naturally-occurring small proteins, are mostly limited to interactions with pre-existing cellular machineries or regulatory pathways rather than *bona fide* enzymatic activities. Specifically, our previously isolated *de novo* small proteins confer antibiotic resistance by altering cell permeability via direct interactions with the cell membrane (Knopp et al. 2019) or by activating a sensor kinase via protein-protein interactions (Knopp et al. 2021). Along similar lines, an earlier study by Digianantonio and Hecht using structurally-constrained and partially randomized DNA libraries showed that selected proteins 102 amino acids in length can rescue an *E. coli* auxotroph caused by the deletion of *serB* (Digianantonio and Hecht 2016). While the precise mechanism of these semi-random proteins has not been elucidated, the dependence of the growth restoration on the deattenuation/increased transcription of the *his* operon and subsequent upregulation of the multi-copy suppressor, HisB, points toward a possible protein-RNA regulatory interaction as the underlying molecular basis of the rescue.

The abundance and mechanisms of RNA regulatory elements, especially those that regulate gene expression in response to direct protein binding, such as ribosomal protein leaders (Fu et al. 2013; Zengel and Lindahl 1994) and Rho-dependent transcription terminators (Banerjee et al. 2006), render them promising potential targets for *de novo* small protein functionality. Additionally, the rescue of auxotrophies is a convenient means to probe for *de novo* small proteins with novel regulatory interactions, as a number of biosynthetic operons are controlled by combinations of different regulatory proteins, RNA elements, and/or small molecules, and many auxotrophic phenotypes can often be suppressed by modulating the expression of alternate enzymes with moonlighting activities (Patrick et al. 2007).

To experimentally investigate the extent to which completely random, unconstrained, and/or noncoding sequences can serve as substrates for natural *de novo* gene evolution, we utilized random sequence expression libraries (Knopp et al. 2019) to select for *de novo* small proteins that can restore the growth of an auxotrophic *E. coli* strain lacking the conditionally essential enzyme, SerB, and characterized the mechanisms responsible for the rescue phenotype. We isolated three small proteins from our screen that are less than or equal to 50 amino acids in length and are novel and distinct from those isolated from past studies. Our selected proteins confer their rescue effect by upregulating the alternative enzyme HisB, and the increase in *hisB* expression is likely caused by direct RNA-binding interactions with the regulatory 5’ end of the *his* operon mRNA transcript. In addition to their small size and gene regulatory roles, the recovered proteins exhibit other traits characteristic of most known naturally-occurring small proteins, providing additional *in vivo* evidence that sORFs encoding novel beneficial functions can indeed originate *de novo* from previously noncoding and/or nonfunctional DNA sequences, without any pre-existing structural or functional scaffolds. These findings add nucleic acids to the list of interacting partners for the novel *de novo* small proteins isolated from *in vivo* random sequence library screens.

## RESULTS

### Selection of novel sORFs that restore growth of an auxotrophic *E. coli* mutant

We used a set of highly diverse expression vector libraries (Knopp et al. 2019) to screen for novel sORFs that confer rescue of the *serB* deletion mutant and other auxotrophic *E. coli* strains. The five libraries encode small proteins ranging from 10 to 50 amino acids in length (Fig. 1A). Three of the libraries (rnd10, rnd20, rnd50a) encode repeats of NNB, where N encodes A, C, G, or T at equal ratios and B encodes only C, G, or T. This restriction was chosen to remove two of the three stop codons to increase the likelihood of obtaining full-length proteins within the sORFs, while maintaining amino acid ratios comparable to that of NNN repeats. Libraries 50b and 50c were biased to encode a higher fraction of primordial and hydrophilic residues, in an effort to recapitulate potential coding sequences in early life (Ring et al. 1972) or generate proteins without a strong hydrophobic core that are structurally flexible (Dunker et al. 2005; Dyson 2016), respectively. The inserts were expressed from an IPTG-inducible low-copy plasmid (an in-house construct denoted as pRD2 containing a p15A origin of replication). A strong ribosome-binding site, a start codon, and three stop codons (one per frame) ensured translational initiation and termination of the randomly generated inserts upon IPTG induction. The total diversity of the libraries was approximately 5.8 × 10^8^.

**Figure 1:**
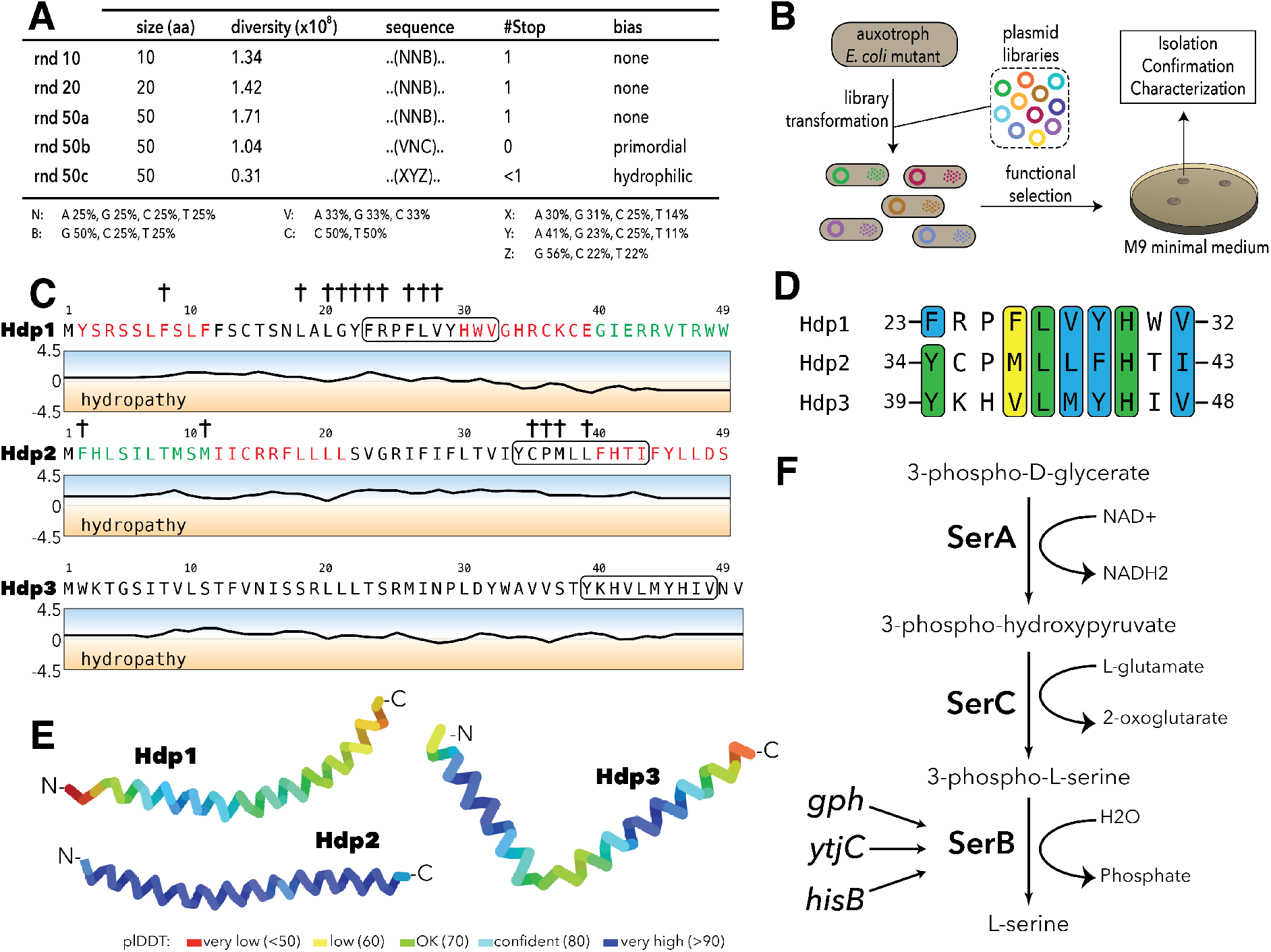
Experimental setup and sequence characteristics of the isolated small proteins. (A) Libraries cloned into the expression vector pRD2. (B) Plasmid transformation into auxotrophic mutants and selection for rescue of auxotrophic mutants. (C) Hydropathy profiles of the three isolated small proteins (Hdp1-3). Colored amino acids denote residues that could (green) or could not (red) be removed while maintaining functionality. Loss of function mutations for Hdp1 and Hdp2 are indicated by †. The box spanning 10 amino acids denotes a region of similarity between the small proteins. (D) Sequence alignment showing the region of similarity shared between the three small proteins. Green coloring indicates identical amino acids, while blue and yellow coloring indicate strongly and weakly similar amino acids, respectively. (E) Structure prediction of Hdp1-3 obtained from AlphaFold (Jumper et al. 2021; Mirdita, Ovchinnikov, and Steinegger 2021). Colors represent the per-residue prediction confidence level (pLDDT), based on the lDDT-cα metric (Mariani et al. 2013). (F) The three enzymes SerA, SerC, and SerB catalyze the last step in L-serine biosynthesis. Essentiality of SerB on minimal medium can be suppressed by overexpression of the phosphatases Gph, YtjC, or HisB.

We used these libraries to select for plasmid clones that could restore growth of the Δ*serB* auxotroph and other auxotrophic *E. coli* mutants on minimal medium. The strains used were single-gene knockout mutants from the KEIO-collection (Baba et al. 2006). A previous study showed that 155 KEIO-strains are unable to grow on M9 minimal medium (Baba et al. 2006; Joyce et al. 2006; Patrick et al. 2007); we selected 74 of these strains for our screen (Table S1.). These mutants included: strict or “tight” auxotrophs that do not form colonies or residual growth on plates even after 2 weeks of incubation, (ii) “leaky” auxotrophs that are able to grow after an extended incubation time, and (iii) auxotrophs with high reversion frequencies, which are generally not able to grow even after prolonged incubation periods, but have a higher frequency of reversion via chromosomal mutations. We transformed all five expression libraries as well as empty vector controls into these 74 auxotrophic mutants and selected three variants that enabled growth on M9 minimal medium (Fig. 1B). While multi-copy suppression of these auxotrophies has previously been shown to be common (Patrick et al. 2007), we were only able to isolate randomly generated sORFs that could rescue the Δ*serB* auxotroph from our screens. The inserts encoded in the three selected variants were designated *his deattenuating protein* 1-3 (Hdp1-Hdp3).

### Small proteins that confer rescue are alpha-helical and share a common sequence motif

To determine if the rescue of the *serB* deletion mutant is facilitated by expression of the mRNA or the encoded protein, we constructed Hdp1 and Hdp2 variants containing frameshifts and premature stop codons. None of these constructs were able to restore growth of the auxotrophic mutant on minimal medium, indicating that the translated proteins are responsible for the rescue. Furthermore, we generated variants encoding the proteins using alternative codons, which extensively changes the nucleotide sequence while maintaining the amino acid sequence. Recoded Hdp2 retained function, demonstrating that the rescue is indeed mediated by the translated sORF. However, recoding Hdp1 resulted in a loss of function, likely due to effects on expression levels and mRNA stability. Therefore, to further demonstrate that the functionality of this insert is also dependent on the encoded protein rather than the mRNA transcript, we constructed an additional Hdp1 variant in which the start codon was removed. This variant did not allow growth on minimal medium, confirming that protein expression is essential for rescue.

The encoded small proteins are hydrophobic 49- and 50-mers (Fig. 1C) and truncation experiments showed that 10 amino acids could be removed from the C- and N-terminus of Hdp1 and Hdp2, respectively, with maintained functionality. Multiple sequence alignments using Clustal Omega (Chojnacki et al. 2017) revealed a motif of 10 amino acids that was shared among all three proteins (Fig. 1D). This motif is predicted to be mainly alpha-helical (Jumper et al. 2021; Mirdita et al. 2021), has not been described previously, and is not recognized in the Pfam database of protein families (El-Gebali et al. 2019) (Fig. 1E). To examine the potential functional role of this similarity region and define other functional regions of the proteins, we performed random-mutagenesis of the *hdp1* and *hdp2* genes and screened for mutant variants that were unable to rescue the Δ*serB* mutant on minimal medium. A majority of the loss-of-function mutations observed were clustered in or near the similarity region, supporting the hypothesis that it has a role in rescue (Fig. 1C).

### Rescue of the Δ*serB* auxotrophy in *E. coli* K12 is *hisB*-dependent

SerB is a phosphatase that catalyzes the conversion of 3-phospho-L-serine to L-serine in the final step of L-serine biosynthesis (Ravnikar and Somerville 1987). To test whether the isolated proteins can bypass the normal pathway for L-serine biosynthesis, we examined if they could rescue auxotrophies caused by the deletion of the enzymes upstream in this linear pathway (Fig. 1F). On minimal medium, none of the Hdp proteins enabled growth of either a Δ*serA* or Δ*serC* mutant, which encode a dehydrogenase and an aminotransferase, respectively, indicating that the Hdps do not re-route the metabolism to synthesize serine via a different pathway, but rather they relieve the need for SerB.

*E. coli* encodes 23 cytoplasmic haloacid dehydrogenase (HAD)-like hydrolases (including SerB), which share limited sequence similarity but have strongly overlapping substrate specificities (Kuznetsova et al. 2006). To determine whether expression of the selected small proteins could functionally replace any of the other HAD-like phosphatases, we tested the growth defects of individual HAD-knockout mutants. Besides Δ*serB*, only Δ*hisB* exhibited an auxotrophic phenotype. However, only the Δ*serB* auxotrophic mutant could be rescued by expression of the isolated proteins.

Past studies showed that the overexpression of two HAD-like phosphatases, HisB and Gph, as well as the non-related putative phosphatase YtjC, can rescue Δ*serB* auxotrophy (Patrick et al. 2007; Yip and Matsumura 2013). We therefore tested if the small protein-mediated rescue is dependent on the presence of any of these three proteins. While removal of YtjC and Gph did not affect the rescue, deletion of *hisB* abolished the ability of Hdp1-Hdp3 to rescue the Δ*serB* auxotrophy. Based on these findings, we hypothesized that the Hdps rescue the lack of SerB by upregulating expression of HisB, which can functionally replace SerB and thereby restore L-serine biosynthesis and growth on minimal medium (Fig. 1F). This is in agreement with the previous Digianantonio and Hecht study which showed that sequence libraries encoding semi-random proteins that fold into pre-defined four-helix bundles could be used to select proteins that upregulate HisB to rescue SerB-deficiency (Digianantonio and Hecht 2016; Fisher et al. 2011).

### The *his* operator region is required for the Hdp-mediated rescue of Δ*serB* auxotropy

Expression of the *his* operon, consisting of the structural genes *hisG, hisD, hisC, hisB, hisH, hisA, hisF* and *hisI*, is regulated by a transcriptional attenuator located near the 5’ end of the *his* operon mRNA transcript that responds to levels of charged tRNA^His^ (Fig. 2A). Under histidine-rich conditions, a histidine-rich leader peptide (HisL) is synthesized and a terminator hairpin forms, resulting in transcription termination. However, under histidine-poor conditions, the ribosome will stall at the HisL histidine codons due to the lack of charged tRNA^His^. As a result, an anti-terminator hairpin forms that allows continued transcription and subsequent expression of the downstream structural genes (Artz and Broach 1975; Blasi and Bruni 1981; Johnston et al. 1980; Kasai 1974). Additionally, the operon promoter is activated upon induction of the stringent response, offering various potential targets with which the Hdps could interact. For simplicity, we designated the aforementioned regulatory region of the *his* operon as the “*his* operator (*his*_operator_),” comprised of the *hisL* leader peptide gene and the transcription attenuator and under the control of the P_*hisL*_ promoter (Fig. 2A).

**Figure 2:**
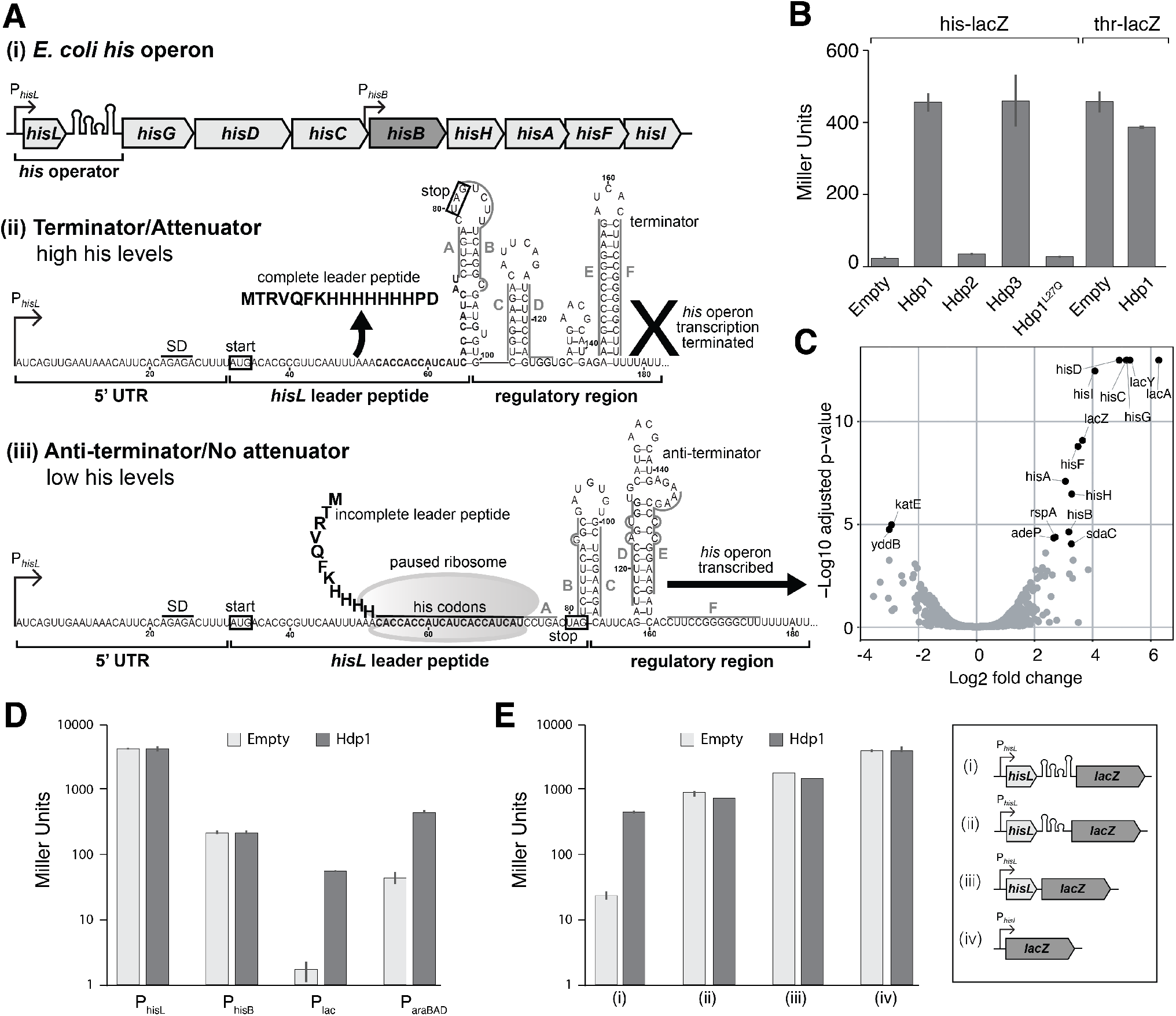
Overview of *his* operon regulation in *E. coli* and regulatory activity of the Hdps. (A) (i) The control region and structural genes of the *his* operon, including *hisB* (highlighted in dark grey), (ii) the *his* operator and RNA secondary structures under histidine-rich conditions, and (iii) the *his* operator and RNA secondary structures under histidine starvation. P*hisL* is the promoter, *hisL* is the leader peptide, SD is the Shine-Dalgarno sequence for *hisL*, the start and stop codons of the *hisL* coding region are boxed. Nucleotides are numbered from the transcription start site, +1. RNA secondary structures are adapted from (Johnston et al. 1980; Kolter and Yanofsky 1982). (B) β-galactosidase activity (in Miller Units) of the strain carrying the full-length P*hisL*-*his*operator-*lacZ* reporter upon expression of Hdp1-3 and the Hdp1 L27Q mutant, and the strain containing a P*thrL*-*thr*operator-*lacZ* reporter upon the expression of Hdp1. (C) “Volcano plot” showing changes in protein expression in the Δ*serB* mutant containing the full-length P*hisL*-*his*operator-*lacZ* reporter upon expression of Hdp1 versus the empty plasmid control. (D) β-galactosidase activity of various *lacZ* reporter constructs under the control of different promoters upon expression of Hdp1 versus the empty plasmid control: *lacZ* transcriptional fusions under the control of the native *hisL* and *hisB* promoters (no *his* operator regulatory sequence) and the full-length *his*operator-*lacZ* reporter construct under the control of the IPTG-inducible P*lac* or arabinose-inducible P*araBAD* promoters. (E) β-galactosidase activity of various truncated P*hisL*-*his*operator-*lacZ* reporter constructs upon expression of Hdp1 versus the empty plasmid control. Refer to Table S3 for the nucleotide sequences of each *lacZ* reporter construct. For the β-galactosidase data presented in panels B/D/E, the values reported represent the mean of three or more independent biological replicates; error bars represent the standard deviation.

To test if increased expression of HisB by the Hdps is dependent on the promoters and/or regulatory region of the *his* operon transcript, we constructed various transcriptional fusions of the *his* operator to the native *lacZ* gene on the chromosome of the Δ*serB* strain to monitor activity *in vivo*. Hdp1 and Hdp3 expression caused an almost 20-fold increase in β-galactosidase activity of the reporter strain carrying the full-length P_*hisL*_-*his*_operator_-*lacZ* fusion (under the control of the native P_*hisL*_ promoter) (Fig. 2B). Only a 2-fold increase was observed for Hdp2, consistent with our observations of Δ*serB* strain growth on minimal medium where cells expressing Hdp1 and Hdp3 grew faster than those expressing Hdp2. Correspondingly, a nonfunctional Hdp1 variant containing a L27Q substitution within the similarity region that was derived from the random mutagenesis screens (i.e. did not rescue the Δ*serB* mutant on minimal medium) did not increase the β-galactosidase activity of the P_*hisL*_-*his*_operator_-*lacZ* fusion reporter strain. To examine the specificity of the Hdp mechanism of action, we also assayed the β-galactosidase activity of a reporter strain carrying the full-length *thr* operator sequence transcriptionally fused to the *lacZ* gene on the chromosome. In *E. coli*, the *thr* biosynthetic operon is regulated by an operator region that is similar in length, structure, and mechanism to that of the *his* operon (Kolter and Yanofsky 1982). As expected, Hdp1 expression had little impact on the β- galactosidase activity of the P_*thrL*_-*thr*_operator_-*lacZ* reporter strain (Fig. 2B). Overall, these data confirm that the Hdps rescue *serB* auxotrophy by increasing *hisB* expression through altering the regulation of the *his* operon via the *his* operator region. Due to its robust activity *in vivo* and the clear negative effect of the L27Q substitution, Hdp1 was used for all subsequent characterization studies.

We next examined if the stringent response is involved in the mechanism of the Hdps, as this pathway is a known regulator of many biosynthetic operons, including the *his* and *trp* operons (Paul, Berkmen, and Gourse 2005; Riggs et al. 1986; Stephens, Artz, and Ames 1975). In addition to the negligible impact Hdp1 expression had on the β-galactosidase activity of the full-length P_*thrL*_-*thr*_operator_- *lacZ* reporter strain, Hdp1 expression did not affect the β-galactosidase activity of a *lacZ* transcriptional fusion under the control of the *rrnB* P1 promoter and accompanying regulatory sequence, which is also regulated by the stringent response (Aseev, Koledinskaya, and Boni 2014; Paul et al. 2004, 2005) (Fig. S1A). Moreover, the rescue effect of Hdp1 was not affected by the deletion of the stringent response genes *relA* and *dksA* in the P_*hisL*_-*his*_operator_-*lacZ* fusion reporter strain (Paul et al. 2004; Turnbull et al. 2019) (Fig. S1B). To further evaluate if Hdp1 expression activates cellular stress response mechanisms, we also performed a global proteome analysis of cells expressing Hdp1 versus those with an empty plasmid control. We used the Δ*serB* strain carrying the full-length P_*hisL*_-*his*_operator_-*lacZ* fusion as the test strain (DA57390, Table S3). The abundances of only 16 proteins were significantly altered upon expression of Hdp1, and 12 of the 14 proteins demonstrating a significant increase in abundance were under the regulation of the *his* operator (i.e. the native *his* operon and modified *lac* operon) (Fig. 2C). These data suggest that Hdp1 does not induce *his* operon expression through the activation or upregulation of stress response pathways and that Hdp1 has a high specificity in its action and does not cause global alterations in gene expression. Due to their small size, HisL and Hdp1 were not recovered in the proteomics samples.

To assess if the Hdps alter HisB expression through direct interactions with either of the two native *his* operon promoters, we generated *lacZ* transcriptional fusions (with no *his* operator regulatory sequence) under the control of the P_*hisL*_ promoter, which is the primary *his* operon promoter located upstream of the *hisL* leader peptide gene, and the P_*hisB*_ promoter, an internal promoter for the operon located just upstream of the *hisB* gene (Grisolia, Riccio, and Bruni 1983). No increase in β- galactosidase activity was observed upon Hdp1 expression for either reporter strain carrying the P_*hisL*_- *lacZ* or P_*hisB*_-*lacZ* transcriptional fusions (Fig. 2D). However, Hdp1 was still able to increase the β- galactosidase activity of *his*_operator_-*lacZ* transcriptional fusions in which the native P_*hisL*_ promoter was replaced with either the P_*lac*_ promoter or the P_*araBAD*_ promoter (induced upon the addition of IPTG or arabinose, respectively), showing that the Hdp rescue mechanism does not require the native *his* operon P_*hisL*_ and P_*hisB*_ promoters.

Finally, no increase in β-galactosidase activity was observed upon Hdp1 expression in strains carrying truncated versions of the *his* operator fused to *lacZ* (Fig. 2E). An Hdp1-mediated increase in β-galactosidase activity was also not observed (i.e. white colonies on MacConkey agar) for a full-length P_*hisL*_-*his*_operator_-*lacZ* reporter strain containing a missense mutation within the *hisL* leader peptide coding sequence (Q5Stop). Taken together, our data demonstrate that the Hdps require the complete and fully functional *his* operator regulatory region, consisting of the full-length and intact *hisL* gene and transcription attenuator sequence, to exert their rescue effect.

### Hdp1 binds the *his* operator mRNA

Based on our β-galactosidase activity data, we hypothesized that the Hdps likely modulate *his* operon expression through direct interaction with the operator region of the *his* operon mRNA transcript. The original Hdp1 protein exhibited very low solubility in water (< 0.2 mg/ml), initially precluding us from performing *in vitro* binding assays. To circumvent this problem, a functional and more hydrophilic Hdp1 variant with increased solubility in water (> 1 mg/ml), named Hdp1-optimized (Hdp1_opt_), was derived from the mutagenesis experiments (Fig. S2). We also generated an Hdp1_opt_ variant containing the previously described L27Q substitution, which completely abolished Hdp1 activity *in vivo*, to serve as a control for our *in vitro* experiments.

Direct binding of Hdp1_opt_ to the full-length *his* operator RNA was confirmed by electrophoretic mobility shift assays (EMSAs). The *K*_d_ value from a 1:1 binding model was estimated as 0.63±0.13 μM. Consistent with our *in vivo* data, EMSAs performed with the nonfunctional Hdp1_opt_ L27Q variant demonstrated a 5-fold reduction in binding affinity *in vitro* (*K*_d_=3.3±0.13 μM) (Fig. 3A, B). However, while a maximum bound fraction of 1 is expected from the experimental setup, the fitted maximum fraction bound was 1.45±0.13 for the L27Q mutant protein. This value results from a slightly sigmoidal shape of the binding curve, suggesting positive cooperativity. We therefore fitted an equation accounting for positive cooperativity to the data, resulting in *K*_d_ values of 0.54±0.08 μM (Hdp1_opt_) and 1.8±0.26 μM (Hdp1_opt_ L27Q), with Hill coefficients of around 1.5 (Fig. S3). While the number of data points collected precludes any strong conclusion regarding the binding model, the EMSAs demonstrate a direct binding interaction between the Hdp1_opt_ protein and the *his* operator RNA and that Hdp1_opt_ displays a higher affinity for the RNA than the L27Q mutant.

**Figure 3:**
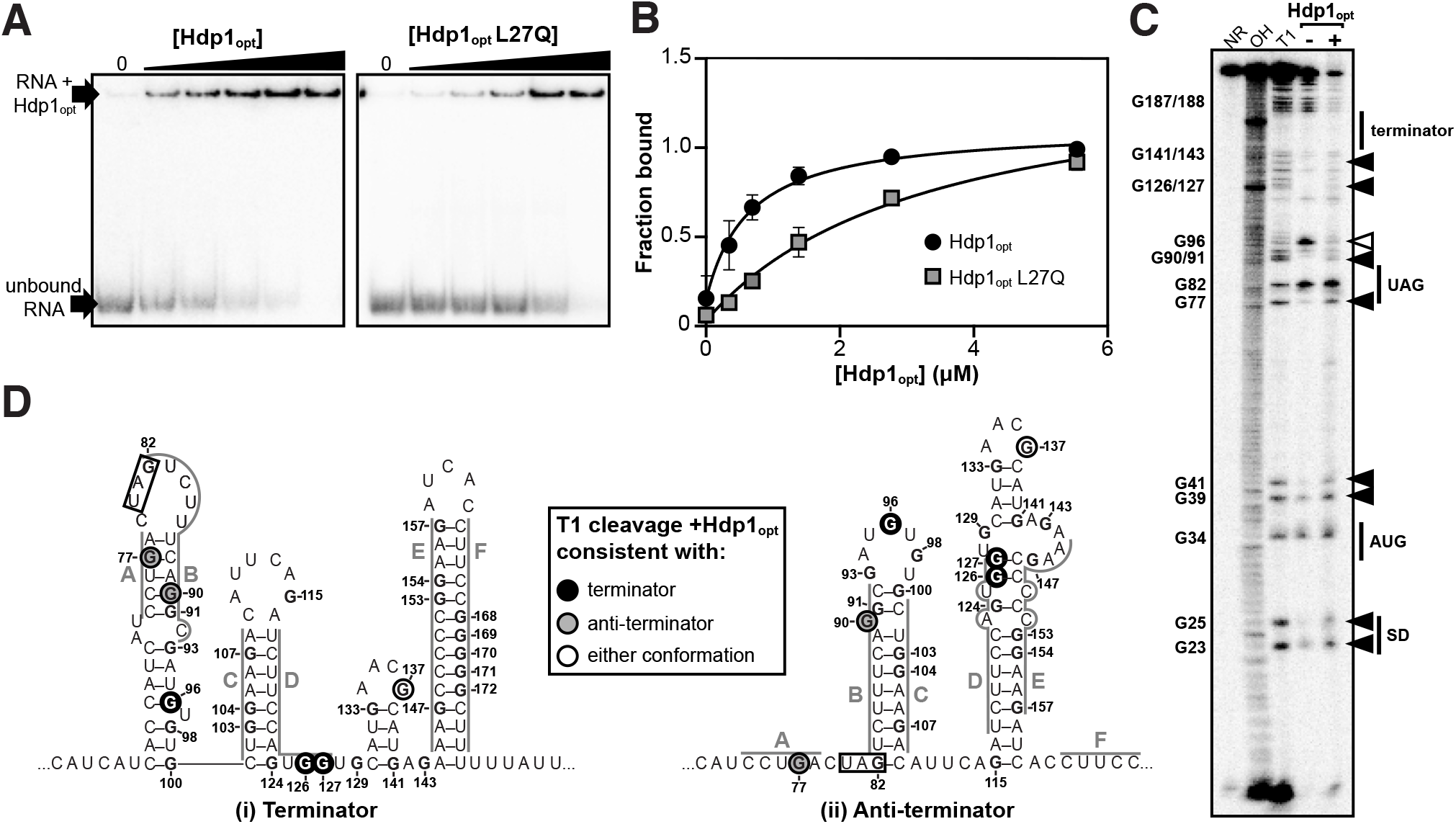
*In vitro* characterization of Hdp1_opt_ binding interactions with the *his* operator mRNA. (A) Electrophoretic mobility shift assays (EMSAs) of the full-length *his* operator RNA in the presence of increasing concentrations of Hdp1opt or the Hdp1opt L27Q mutant (0-5.5 μM). EMSAs were repeated independently three or more times for each protein with similar results; representative gels are shown. (B) Binding curves obtained from fitting the data quantified from the EMSAs to a standard 1:1 hyperbolic binding model, with *K*d values of 0.63±0.13 μM for Hdp1opt and 3.3±0.13 μM Hdp1opt L27Q, and maximum fraction bound values of 0.97±0.06 for Hdp1opt and 1.45±0.13 for Hdp1opt L27Q. Each data point represents the mean of three or more independent experimental replicates; error bars represent the standard error. Some error bars fall within the boundaries of the markers. For the parameters used and the EMSA data fit to a binding model accounting for positive cooperativity, refer to Fig. S3. (C) RNase T1 probing gel for the full-length *his* operator RNA in the absence and presence of Hdp1opt (0 and 5.5 μM, respectively). NR denotes RNA subject to no reaction, OH indicates partial alkaline hydrolysis, and T1 is an RNase T1 digest of the RNA under denaturing conditions used to map the RNA sequence. Numbering of G nucleotides is shown on the left. Arrows highlight changes in RNA cleavage in the presence of Hdp1opt: black arrows indicate nucleotides with increased cleavage and white arrows indicate reduced cleavage. Sequence and/or structure characteristics of the *his* operator RNA are also indicated on the right (i.e. the Shine-Dalgarno sequence (SD), start codon (AUG), and stop codon (UAG) of the *hisL* leader peptide coding sequence). This experiment was repeated three independent times with similar results; a representative gel is shown. (D) Nucleotides demonstrating a change in RNase T1 cleavage in the presence of Hdp1opt mapped to both the terminator (i) and anti-terminator (ii) conformations of the *his* operator RNA. The conformation consistent with cleavage in the presence of Hdp1opt is color-coded according to the legend; the *hisL* stop codon is boxed. Nucleotides are numbered from the transcription start site, +1.

### Hdp1 binding alters *his* operator RNA secondary structure

To assess if portions of the full-length *his* operator RNA undergo conformational changes upon Hdp1_opt_ binding and/or which regions are necessary for Hdp1_opt_ interaction, we performed nuclease protection assays with RNase T1 in the presence and absence of Hdp1_opt_ (±5.5 μM Hdp1_opt_, the maximum concentration used in the EMSAs). RNase T1 specifically cleaves RNA at single-stranded or unprotected G nucleotides. The *his* operator RNA can adopt two mutually exclusive conformations (Johnston et al. 1980; Kolter and Yanofsky 1982), and probing the RNA structure in the absence of Hdp1_opt_ showed hallmarks of both conformations (Fig. 3C, D). For instance, protection of cleavage at G77 and G90 is consistent with the terminator conformation, while the strong cleavage at G96 and protection at G126-127 is consistent with the anti-terminator structure, suggesting a mixture of the two conformations, or possibly alternative conformations, in the RNA sample.

Several regions of the *his* operator RNA demonstrated increased cleavage in the presence of Hdp1_opt_ and the cleavage pattern suggests a transition between the terminator and the anti-terminator conformations. Specifically, while the increased cleavage of nucleotides G126 and G127 in the presence of Hdp1_opt_ is consistent with the terminator conformation of the *his* operator RNA, the increased cleavage of nucleotides G77 and G90 is consistent with the anti-terminator structure (Fig. 3C, D). Interestingly, only one nucleotide, G96, exhibited reduced cleavage upon the addition of Hdp1_opt_. The protection of G96 in the presence of the protein is consistent with the terminator conformation of the *his* operator RNA, however, it may also indicate a region of the RNA that is directly shielded by protein binding.

The intensity of T1-cleavage products corresponding to the *hisL* Shine-Dalgarno sequence and start codon (nucleotides G23-G41) also increased in the Hdp1_opt_-bound RNA. In addition to the impact the Hdps have on *his* operon transcription, it is also possible that the 5’-region of the RNA becomes more accessible to the ribosome in the presence of Hdp1_opt_, thus facilitating more efficient translation of the *his* operon and further increasing HisB expression. However, the biological relevance of this T1-cleavage pattern is difficult to discern, as the 5’-region of the *his* operator RNA is predicted to remain relatively unstructured in either conformation (Johnston et al. 1980; Kolter and Yanofsky 1982).

Some highly structured regions could not be fully resolved in neither the T1 probing assays nor the denaturing T1 reference ladder, particularly the residues that comprise the terminator stem (E+F, nucleotides G153-G171) (Fig. 3C). This indicates that the G-C-rich stem is difficult to disrupt, even under denaturing conditions and with a high RNase T1 concentration (Johnston et al. 1980; Kolter and Yanofsky 1982). Similarly, the region corresponding to C in Fig. 3D (nucleotides G103-G107) is also difficult to resolve in both the T1 ladder and the probing assays. This region is base-paired in both *his* operator RNA conformations, thus it may also be difficult to disrupt and subsequently generate corresponding cleavage products.

Despite these limitations and although we were unable to identify any specific Hdp1_opt_ binding site, the T1 probing assays do provide insight into the general regions of the *his* operator RNA that are impacted by protein binding. Possibly, Hdp1_opt_ binding results in changes to the *his* operator RNA secondary structure that favor the canonical anti-terminator conformation or promote the formation of an alternative anti-terminator structure. Sufficient stabilization of an anti-terminator conformation and/or destabilization of the *his* operator terminator upon Hdp binding would likely allow increased expression of *hisB* and ultimately the growth of the Δ*serB* auxotrophic mutant on minimal medium.

## DISCUSSION

The recent identification of a multitude of small proteins encoded by sORFs raises intriguing questions about their functional roles and evolutionary origins. With regard to their emergence, one possible mechanism for generating sORFs is by degradation of larger genes into pseudogenes containing shorter coding regions (Hemm et al. 2008). Also in *E. coli*, other sORFs have been identified in prophage regions and thus may have origins stemming from an ancestral phage genomic integration event (Hemm et al. 2008; VanOrsdel et al. 2018; Weaver et al. 2019). However, these mechanisms only address the origins of a fraction of small protein-coding genes. A large majority of sORFs identified to date demonstrate little to no conservation between closely related phylogenetic classes and lack the genomic context to support or indicate either of the aforementioned mechanisms, suggesting that many of these small genes may have emerged *de novo* from previously non-genic sequences. Nonetheless, while previous studies show that novel functions, including variants with enzymatic activity, can be selected in bacteria from templates featuring localized randomization within larger functional progenitor sequences and/or structural scaffolds (Digianantonio and Hecht 2016; Hoegler and Hecht 2016; Smith, Mularz, and Hecht 2015), there is limited experimental evidence demonstrating that *de novo* genes can emerge from the expression of completely random and/or nonfunctional DNA (Knopp et al. 2019, 2021). In this work, we identified *de novo* small proteins with RNA-binding regulatory activities.

We isolated three independent small proteins (Hdp1-Hdp3) that rescued an *E. coli* Δ*serB* mutant by upregulating the expression of an alternate enzyme, HisB, and for Hdp1 we showed direct binding interactions with the 5’ end of the *his* operon mRNA transcript. The *his* operon is regulated by a transcriptional attenuation mechanism (Blasi and Bruni 1981; Johnston et al. 1980) as well as the stringent response that acts on the promoter and transcription initiation (Riggs et al. 1986; Stephens et al. 1975). Analysis of the β-galactosidase activity of several *his*_operator_-*lacZ* transcriptional fusions in response to the presence of Hdp1 showed that: (i) Hdp1 acts at the level of transcriptional attenuation (rather than the promoter), (ii) the stringent response is not involved in the Hdp1-mediated effects on *his* operon expression, and (iii) Hdp1 action requires the intact *his* operator region.

In addition, *in vitro* experiments demonstrated that Hdp1 directly binds to the *his* operator RNA and that protein binding results in changes to the RNA secondary structure that likely allow increased transcription of the *his* operon, including *hisB*. Using a more soluble variant of Hdp1 (Hdp1_opt_) that maintained the Δ*serB*-rescue phenotype, we showed by EMSAs that Hdp1_opt_ binds to the full-length *his* operator RNA with micromolar affinity and that the inactive protein mutant, Hdp1_opt_ L27Q, has a 5-fold reduction in affinity. Most RNA-protein regulatory interactions exhibit *in vitro K*_d_ values within the nanomolar range or lower (Ryder, Recht, and Williamson 2008). However, the moderate binding affinity of Hdp1_opt_ to the *his* operator RNA is not surprising, as the protein was directly selected from a random library without any further optimization.

More in-depth characterization of the binding interaction between Hdp1_opt_ and the *his* operator RNA could be complicated by the possibility that the Hdps may exist as multimers rather than monomeric proteins, as alpha-helical proteins with predominantly hydrophobic residues often have a propensity to oligomerize (Li, Wimley, and Hristova 2012). Hdp multimerization is conceivable, as the semi-random proteins isolated from the previous Digianantonio and Hecht study that likely rescue Δ*serB* auxotrophy via a similar (but undetermined) mechanism of action were designed to fold into four-helix structures (Digianantonio and Hecht 2016). Albeit beyond the scope of this work, further investigations are needed to determine the precise molecular mechanisms underlying the binding model and/or stoichiometry of the Hdp-*his* RNA interaction.

As mentioned above, Digianantonio and Hecht (Digianantonio and Hecht 2016) also recovered several proteins (SynserB1-4) that rescued growth of a Δ*serB* auxotrophic mutant via HisB upregulation by utilizing libraries of partially randomized sequences. Even though the exact mechanism of the SynserB-mediated rescue is not characterized, it is likely that both the SynserBs and Hdps share a similar mechanism, as both upregulate HisB expression in a stringent response-independent manner. Despite the functional similarity, there is no sequence similarity between the SynserBs and Hdps, demonstrating that the selected function is not dependent on a specific extended sequence motif. Interestingly, we observed that expression of Hdp1 causes a specific upregulation of the *his* operon. In contrast, the SynSerB3 protein showed a rather unspecific effect on gene expression, with more than 600 proteins being affected (including most of the amino acid biosynthetic operons). It is possible that the differences in global proteome/transcriptome changes observed for *synserB3* and *hdp1* are due to different experimental conditions. SynSerB3 was tested in a *serB*-deficient mutant in minimal medium, which results in severely reduced growth, while Hdp1 was tested in rich medium, which allowed for wild-type growth rates and comparisons with empty plasmid control strains.

Based on the binary pattern constraining the library design in the Digianantonio and Hecht study, the selected SynserB proteins form four-helix bundles. While this library design increases the chances of recovering well-structured proteins that likely reach high cellular concentrations without aggregating, *bona fide de novo* proteins evolved from completely random sequences do not share this privilege and will mostly form insoluble aggregates and be targeted for degradation (Prijambada et al. 1998). Additionally, using libraries with pre-existing structural properties further limits the selection of novel functionalities to those within the defined sequence and structural constraints. By utilizing expression libraries encoding completely random sORFs, we have shown that a new functionality can be selected truly *de novo*, without any pre-defined structural boundaries, demonstrating that pre-existing structural scaffolds are not necessary for a novel protein to be expressed and achieve biological function with high specificity. However, randomized sequences in nature are likely not completely random, as they may contain remnants of previously optimized structural or functional sequence motifs, which might increase the probability of acquiring a new function. Yet, the use of completely random sequences is a suitable starting point for proof-of-concept demonstration of the *de novo* evolution of small protein-coding genes, as they serve as a true null model when screening for functionality, and unlike pre-existing genomic sequences, they introduce little implicit bias or constraints during the selection process.

Given that the majority of small proteins described to date are predominately composed of hydrophobic amino acids (Hemm et al. 2008; Vakirlis et al. 2020), it is not surprising that all functional small proteins we have selected are also hydrophobic. As our previously recovered antibiotic resistance-conferring proteins are localized to the membrane, the strong hydrophobicity is linked to their functionality. However, despite the transmembrane domain predictions, it is unlikely that the Hdps exclusively associate with the cell membrane, as their mechanism involves regulating transcription and the solubility-optimized variant of Hdp1 (Hdp1_opt_) is not predicted to target the membrane (Fig. 1C, S2A). Nevertheless, it is interesting to consider that hydrophobicity is an important contributor to functionality in random sequence space. This concept is supported by recent work from Vakirlis *et al*. (Vakirlis et al. 2020), which identified that the beneficial fitness effects of emerging ORFs were found to be associated with the potential to produce transmembrane domains, and that thymine-rich intergenic regions in particular were identified as a reservoir for encoding transmembrane domains.

Hdps, as well as the small proteins selected from our past screens (Knopp et al. 2019, 2021), demonstrate many qualities intrinsic to most known naturally-occurring bacterial small proteins: (i) they are ≤ 50 amino acids in length, (ii) contain a high percentage of hydrophobic amino acids, (iii) are predicted to form predominantly alpha-helical structures, (iv) are predicted to associate with the membrane, and (v) they do not encode an enzymatic activity, but rather act as regulators/interactors to modulate a specific process (Carvunis et al. 2012; Storz et al. 2014). The previously isolated aminoglycoside resistance-conferring proteins (Arps) provide resistance by membrane insertion, disruption of the proton motive force, and a subsequent reduction in antibiotic uptake (Knopp et al. 2019), while the colistin resistance-conferring proteins (Dcrs) exert their effect via direct protein-protein interactions which activate the sensor kinase PmrB, resulting in Lipid A modifications and decreased affinity towards colistin (Knopp et al. 2021). This study extends the previously known spectrum of interacting partners of experimentally selected *de novo* small proteins to include nucleic acids, and exemplifies how direct binding of a *de novo* protein to an RNA regulatory element can upregulate expression of a biosynthetic operon and restore growth of an auxotrophic *E. coli* strain. Directed evolution of our selected *de novo* small proteins could shed further light on the evolutionary constrains governing the emergence of new genes. For example, will selection fine-tune the function, expression, and/or stability of these proteins, and will these evolved variants converge to pre-existing genes or differentiate to fill a novel and distinct structural and/or functional niche not yet occupied by naturally-occurring genes?

## MATERIALS AND METHODS

### Strains

All strains were derivatives of *Escherichia coli* MG1655 (F^-^, lambda^-^, *rph-1*). Gene deletion mutants were constructed by P1 transduction from the corresponding KEIO collection strains (derivatives of *E. coli* K-12 BW25113: F^-^, DEL(*araD-araB*)567, *lacZ4787*(del)::*rrnB-3*, LAM^-^, *rph-1*, DEL(*rhaD-rhaB*)568, *hsdR514*) (Baba et al. 2006). All *lacZ* gene fusions were constructed using *E. coli* NM580 and transferred to the test strains by P1 transduction (Battesti, Majdalani, and Gottesman 2015). For all strains generated in this study, aliquots (1 ml) from overnight cultures were cryopreserved with 10% DMSO and stored at -80°C. Where specified, strains were grown in/on lysogeny broth (LB Miller; 10 g/L NaCl, 10 g/L tryptone, 5 g/L yeast extract; Sigma Aldrich), LB supplemented with 1.5% (w/v) agar (LA), Mueller-Hinton broth (MH; 17.5 g/L casein acid hydrolysate, 3 g/L beef extract, 1.5 g/L starch; BD Difco), or MH supplemented with 1.5% (w/v) agar (MHA).

### Library construction

Expression vector libraries encoding 10, 20 and 50 amino acid-long sORFs were constructed as previously described (Knopp et al. 2019). Briefly, randomized oligonucleotides were complemented by primer extension and ligated into the expression vector pRD2 using BamHI and PstI restriction sites. Ligations were transformed into electrocompetent NEB5-alpha *E. coli* (New England Biolabs) and plated on LA plates containing 100 µg/ml ampicillin. After overnight incubation at 37°C, the cells were collected from the plate and plasmid pools were extracted using the NucleoBond Xtra Midi Kit (Macherey-Nagel) according to the manufacturer’s recommendation.

### Selection of functional proteins

We selected 74 KEIO strains that were previously described to exhibit auxotrophic growth behavior (Baba et al. 2006; Joyce et al. 2006) including strains with a strong and weak (leaky) auxotrophic phenotype (see Table S1 for a complete list of the tested strains). An overnight culture of each single-gene knockout auxotroph was diluted 1:200 in 100 ml pre-warmed LB medium and incubated at 37°C and shaking at 200 rpm. When the cultures reached the target OD_600_ of 0.2, flasks were quickly cooled in ice water for 10 minutes. The cells were pelleted by centrifugation at 4500xg and 4°C and washed three times with cold 10% glycerol. Finally, cell pellets were resuspended in 300 µl 10% glycerol. Cells (40 µl) were mixed with 2 µl of each plasmid library (or empty vector control) in a chilled microcentrifuge tube and incubated on ice for 5 minutes. The mixture was transferred to an electroporation cuvette with a 1 mm gap width and transformed using a Gene Pulser Xcell electroporator (BioRad) with 1.8 kV, 400 Ω, and 25 µF settings. The cells were recovered in 1 ml SOC medium (20 g/L tryptone, 5 g/L yeast extract, 0.5 g/L NaCl, 10 mM MgCl_2_, 0.25 mM KCl, and 4 g/L glucose) for 1.5 hours at 37°C and shaking at 200 rpm. After recovery, the cells were washed twice with phosphate-buffered saline (PBS; 8 g/L NaCl, 0.2 g/L KCl, 1.44 g/L Na_2_HPO_4_, and 0.24 g/L KH_2_PO_4_) and resuspended in 1 ml PBS. An aliquot of each transformation (10 µl) was subjected to a dilution series, plated on LA supplemented with 100 µg/ml ampicillin, and incubated overnight at 37°C to determine transformation efficiencies. The remaining 990 µl were pelleted, washed twice with 1 ml PBS, resuspended in 200 µl PBS, and spread on M9 minimal medium plates (6 g/L Na_2_HPO_4_, 3 g/L KH_2_PO_4_, 0.5 g/L NaCl, 1 g/L NH_4_Cl, 1 mM MgSO_4_, 0.1 mM CaCl_2_, 1.5% (w/v) agar) containing 0.2% glucose, 1 mM IPTG, and 100 µg/ml ampicillin. The plates were incubated for up to 14 days at 37°C and regularly inspected for growth of colonies.

When the empty vector control transformations showed growth of a faint lawn, plates containing this auxotrophic strain were discarded. Colonies that appeared during the selection were re-streaked on M9 minimal medium plates containing 0.2% glucose, 1 mM IPTG, and 100 µg/ml ampicillin, and the original plate was incubated further to allow the growth of additional slower-growing colonies. Plasmids were isolated from overnight cultures inoculated with the re-streaked colonies using the EZNA Plasmid Mini Kit (Omega Bio-Tek) and re-introduced into the parental plasmid-free auxotrophic strain to confirm whether the rescue was plasmid-mediated or due to chromosomal mutations. Plasmids that were confirmed to mediate rescue of the auxotrophic *E. coli* mutant were sequenced, and the insert was re-synthesized and cloned into the empty pRD2 vector to confirm that the insert causes the rescue rather than any alterations on the plasmid. In the cases where all experiments indicated a rescue mediated by the insert, further mechanistic characterizations were conducted.

### Plasmid transformation

For transformation or re-transformation of miniprepped plasmids, 2 ml of an overnight culture were washed three times with 10% glycerol and the pellet was resuspended in 200 µl 10% glycerol. Cells (40 µl) were mixed with 200 ng of the desired plasmid and incubated on ice for 5 minutes. The cell/plasmid mixture was transferred to an electroporation cuvette with 1mm gap and transformed using a Gene Pulser Xcell electroporator (BioRad) with 1.8 kV, 40 0Ω, and 2 µF settings. Cells were recovered in 1 ml pre-warmed MH for 1 hour at 37°C and shaking at 200 rpm. Cells were then plated on MHA with 100 µg/ml ampicillin and incubated overnight at 37°C and select transformants were subsequently purified in an additional re-streak.

### Sequence analysis

Local sequencing analyses were performed using CLC Main Workbench (Qiagen). Secondary structure predictions were performed using AlphaFold (Jumper et al. 2021; Mirdita et al. 2021). Multiple sequence analyses were performed using Clustal Omega (Sievers et al. 2011) using standard parameters. Hydropathy scores were determined using the integrated CLC tool applying the Kyte-Doolittle hydrophobicity scale and a window size of 11.

### Construction of protein variants

Variants of the isolated inserts were either ordered as ready-made constructs from Geneart (Thermo Fisher) sub-cloned into pRD2 or constructed by cloning of annealed oligos. For the latter, two complementary oligonucleotides (Eurofins Genomics) containing single-stranded ends corresponding to BamHI and PstI cleavage sites were combined with annealing buffer (10 mM Tris pH 7.5, 100 mM NaCl, 100 µM EDTA) at equimolar ratios to a final concentration of 1 µM in a microcentrifuge tube and transferred to 95°C hot water, which was then slowly cooled to room temperature. The resulting double-stranded DNA was then ligated into the BamHI/PstI-digested pRD2 vector using Ready-To-Go T4 DNA Ligase (GE Healthcare Life Sciences) according to the manufacturer’s recommendation. The ligation reaction was purified using the GeneJET Gel Extraction Micro Kit (Thermo Fisher). To transform the final construct, an overnight culture of *E. coli* NEB5-alpha (New England Biolabs) was diluted 1:100 in LB medium and grown to OD_600_ of 0.2. The cells were then cooled on ice for 10 minutes and washed three times with cold 10% glycerol, 40 µl of cells were mixed with 2 µl of each ligation reaction, and transformed using the Gene Pulser Xcell electroporator (BioRad) with 1.8 kV, 400 Ω, and 25 µF settings. Cells were recovered in 1 ml pre-warmed MH for 1 hour at 37°C and shaking at 200 rpm. Transformants were selected on LA plates containing the appropriate antibiotics. Plasmids were isolated from the transformants using the EZNA Plasmid Mini Kit (Omega Bio-Tek) and verified via sequencing. Confirmed constructs were transformed into the strains of interest as described above.

### Site-directed mutagenesis

PCR-amplification of the entire plasmid containing the gene of interest was performed with Phusion DNA polymerase (Thermo Fisher) using two complementary oligonucleotides containing a stretch of overlapping bases with the desired mutation in the middle (Table S2). The reaction product was purified using GeneJET Gel Extraction Kit (Thermo Fisher) and digested with DpnI (Thermo Fisher) for 1 h at 37°C to remove the template DNA. After digestion, the PCR product was purified and transformed into NEB 5-alpha electrocompetent *E. coli* (New England Biolabs) according to the manufacturer’s protocol, plated on LA plates containing 50 µg/ml of ampicillin, and incubated overnight at 37°C. Transformants were inoculated into fresh LB medium supplemented with 50 µg/ml of ampicillin and grown overnight at 37°C with shaking at 200 rpm. Plasmids were purified from the overnight cultures using the EZNA Plasmid Mini Kit (Omega Bio-Tek) according to the manufacturer’s protocol. Presence of the desired nucleotide substitutions were confirmed by Sanger sequencing.

### Random mutagenesis

Random mutagenesis of the *hdp1* and *hdp2* genes was performed with the GeneMorph II Random Mutagenesis Kit (Agilent) using the recommended conditions for obtaining a mean mutation rate of 1 mutation per PCR product. The resulting PCR product was used as a megaprimer for whole-plasmid PCR using Phusion DNA polymerase (Thermo Fisher) (Sarkar and Sommer 1990), which was then purified using the GeneJET PCR Gel Extraction Kit (Thermo Fisher) and digested with DpnI (Thermo Fisher) for 1 h at 37°C to remove the original template plasmid. After digestion, the PCR product was purified again and transformed into electrocompetent DA57390 cells (Table S3), which were plated on MacConkey (BD Difco) plates containing 10% lactose and 50 μg/ml of ampicillin and incubated overnight at 37°C. White colonies indicated loss-of-function mutations in the mutagenized genes. These colonies were inoculated into the fresh LB medium supplemented with 50 μg/ml ampicillin and grown overnight at 37°C with shaking at 200 rpm. The plasmids were purified using the EZNA Plasmid Mini Kit (Omega Bio-Tek) according to the manufacturer’s protocol. The nucleotide substitutions were identified by Sanger sequencing.

### Chromosomal *lacZ* fusions

Chromosomal fusions to the *lacZ* reporter gene were generated using the lambda-red recombination system described in (Battesti et al. 2015). A full list of the *lacZ* fusion genetic constructs and their sequences is shown in Table S3. In this system, *E. coli* strain NM580 carries the temperature-inducible lambda *red* gene, which is used for a standard recombineering technique (Sharan et al. 2009). The sequence of interest replaces the arabinose-inducible *ccdB* toxin gene and adjacent kanamycin-resistance gene upstream of *lacZ*. A zeocin resistance cassette located upstream of the fusion was used for the selection of P1 transductants. *E. coli* strain NM580 was streaked from the frozen stock on LA plates with 1% glucose and 50 μg/ml of kanamycin and grown at 30°C overnight. One colony from the plate was inoculated into 3 ml of fresh LB broth supplemented with 1% glucose and 50 μg/ml kanamycin and grown at 30°C with shaking at 200 rpm until an OD_600_ of 0.2 was reached. The cultures were transferred to a 43°C water bath, incubated for 20 min with shaking, and cooled on ice. Cells were collected and washed three times with 20 ml of ice-cold 10% glycerol. The cells were pelleted and resuspended in 200 μl of ice-cold deionized water. Aliquots (50 μl) were used for electroporation with the corresponding PCR product (in 0.5 μl of water) according to manufacturer’s protocol (BioRad). Cells were recovered in 1 ml of LB with 1% glucose for 1 h at 37°C with shaking at 200 rpm. Cells were then pelleted and plated on LA supplemented with 0.2% arabinose and 20 μg/ml zeocin. After overnight incubation at 37°C, kanamycin-susceptible colonies were confirmed with PCR and Sanger sequencing.

PCR products for the recombineering were designed to have 40 base overlaps with the chromosomal region necessary for recombination and were amplified using Phusion DNA polymerase (Thermo Fisher). When necessary, two-step megaprimer PCR was used, wherein the product synthesized in the first step served as a primer in the second step (Sarkar and Sommer 1990). Oligonucleotides used are listed in Table S2. P1 phage transductions were used to transfer the *lacZ* reporter constructs generated in *E. coli* NM580 to the test strains of interest according to published methods (Battesti et al. 2015).

### β-galactosidase activity assays

Overnight cultures were grown in triplicate in LB broth at 37°C, and 300 μl of each culture was used to inoculate 3 ml of fresh LB medium supplemented with 1 mM IPTG and 50 μg/ml ampicillin. For the arabinose-inducible P_*araBAD*_ reporter strain, 0.2% of arabinose was also added to the medium. Cells were grown to early stationary phase (OD_600_ 1.0-2.2), harvested, and β-galactosidase activity was assayed and measured using a Bioscreen C machine (Growth Curves AB, Finland) essentially as described in (Kacar et al. 2017). Reaction time course was measured at 28°C and 420 nm wavelength at 1 min intervals without shaking and β-galactosidase activity in Miller Units was calculated using the following formula, where t2 and t1 are times of the measurements in the linear range of the reaction: 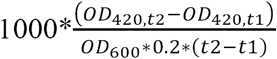 (modified from (Zhang and Bremer 1995)). The values reported represent the mean of three or more independent biological replicates; error bars represent the standard deviation.

### Sample preparation for total proteome analysis

For the proteomic analysis, cells expressing either the empty vector control or *hdp1* were grown until late stationary phase (OD_600_ = 2.5-3) in LB medium in the presence of 1mM IPTG and 50 μg/ml ampicillin. Cells were collected and cell weight was determined. Samples were prepared and an equal amount of each sample was separated on an SDS-PAGE until the dye front reached the bottom of the gel. The whole lane from each sample was cut out from the gel for further analysis. Experiments were performed with two independent biological replicates.

### Label-free proteomic quantification

Each gel lane was cut into six separate pieces, and proteins were reduced in-gel with 10 mM DTT in 25 mM NH_4_HCO_3_, alkylated with 55 mM iodoacetamide in 25 mM NH_4_HCO_3_, and thereafter digested with 17 ng/μl sequencing-grade trypsin (Promega) in 25 mM NH_4_HCO_3_ using a slightly modified in-gel digestion protocol (Shevchenko et al. 1996). The resulting peptides were then eluted from the gel pieces using 1% (v/v) formic acid (FA) in 60% (v/v) acetonitrile, dried down in a vacuum centrifuge (ThermoSavant SPD SpeedVac, Thermo Scientific), and finally dissolved in 1% (v/v) FA.

### Liquid chromatography and mass spectrometry

Peptide samples were desalted using Stage Tips (Thermo Fisher) according to the manufacturer’s protocol, and subsequently dissolved in 0.1% (v/v) FA (solvent A). Desalted samples were separated by RP-HPLC using a Thermo Scientific nLC-1000 with a two-column setup, where an Acclaim PepMap 100 (2 cm x 75 μm, 3 μm particles; Thermo Fisher) pre-column was connected in front of an EASY-Spray PepMap RSLC C18 reversed phase column (50 cm x 75 μm, 2 μm particles; Thermo Fisher). The column was heated to 35°C and equilibrated in solvent A. A gradient of 2–40% solvent B (acetonitrile and 0.1% (v/v) FA) was run at 250 nL/min for 3 h. The eluted peptides were analyzed on a Thermo Scientific Orbitrap Fusion Tribrid mass spectrometer, operated at a Top Speed data-dependent acquisition scan mode, ion-transfer tube temperature of 275°C, and a spray voltage of 2.4 kV. Full scan MS spectra (m/z 400 – 2000) were acquired in profile mode at a resolution of 120,000 at m/z 200, and analyzed in the Orbitrap with an automatic gain control (AGC) target of 2.0e5 and a maximum injection time of 100 ms. Ions with an intensity above 5.0e3 were selected for collision-induced dissociation (CID) fragmentation in the linear ion trap at a collision energy of 30%. The linear ion trap AGC target was set at 1.0e4 with a maximum injection time of 40 ms, and data was collected at centroid mode. Dynamic exclusion was set at 60 s after the first MS1 of the peptide. The system was controlled by Xcalibur software (version 3.0.63.3; Thermo Scientific). Quality control of the instrument was monitored using the Promega 6×5 LC-MS/MS Peptide Reference Mix before and after each MS experiment run, and analyzed using PReMiS software (version 1.0.5.1, Promega).

### Mass spectrometric data analysis

Data analysis of raw files was performed using MaxQuant software (version 1.6.2.3) and the Andromeda search engine (Cox et al. 2009; Tyanova, Temu, and Cox 2016), with cysteine carbamidomethylation as a static modification and methionine oxidation and protein N-terminal acetylation as variable modifications. First search peptide MS1 Orbitrap tolerance was set to 20 ppm and iontrap MS/MS tolerance was set to 0.5 Da. Match between runs was enabled to identify peptides in fractions where only MS1 data were available. Minimum LFQ ratio count was set to 2, and the advanced ratio estimation option was enabled. Peak lists were searched against the UniProtKB/Swiss-Prot *Escherichia coli* K12 proteome database (UP000000625, version 2019-03-27), including the Hdp1 protein sequence, with a maximum of two trypsin miscleavages per peptide. The contaminants database of MaxQuant was also utilized. A decoy search was made against the reversed database, where the peptide and protein false discovery rates were both set to 1%. Only proteins identified with at least two peptides of at least 7 amino acids in length were considered reliable. The peptide output from MaxQuant was filtered by removing reverse database hits, potential contaminants and proteins only identified by site (PTMs). Differential expression analysis was performed by the DEP 1.7.0 package for Bioconductor and R (Zhang et al. 2018). The LFQ intensity data was normalized by the variance stabilizing transformation (vsn) method, and missing values were imputed by a maximum likelihood-based imputation method using the EM algorithm (Gatto and Lilley 2012). Protein-wise linear models and empirical Bayes statistics using LIMMA was used for the differential expression calculation (Ritchie et al. 2015). The *P*-values were adjusted for multiple testing using the Benjamini–Hochberg method (Benjamini and Hochberg 1995). The mass spectrometry proteomics data have been deposited to the ProteomeXchange Consortium (http://proteomecentral.proteomexchange.org) via the PRIDE partner repository (Perez-Riverol et al. 2019) with the dataset identifier PXD014049.

### Electrophoretic mobility shift assays (EMSAs)

The region corresponding to the full-length *his* operator sequence was PCR-amplified from *E. coli* MG1655 genomic DNA using a forward primer that contained the T7 promoter sequence (Table S2). The resulting PCR product was column-purified using the GeneJet Gel Extraction Kit (Thermo Fisher), and RNA was transcribed from this construct (0.2 μg) using the MEGAscript T7 RNA Polymerase Kit (Life Technologies) according to the manufacturer’s protocol, with overnight incubation. The RNA was DNase-treated using the TURBO DNA-free Kit (Invitrogen), and gel purified by 6% denaturing PAGE (Milligan et al. 1987). The purified RNA (60 pmol) was then dephosphorylated using CIAP (Thermo Fisher), phenol-chloroform extracted and ethanol precipitated, and 5 pmol was 5’-end labeled with 20 μCi of γ-P^32^ ATP using Polynucleotide Kinase (Thermo Fisher) and subsequently purified using an Illustra MicroSpin G-50 column (GE Healthcare). The Hdp1_opt_ and Hdp1_opt_ L27Q proteins used for the *in vitro* experiments were synthesized by GenScript and resuspended in water for a stock concentration of 1 mg/ml.

For the binding reactions, a 2-fold dilution series was prepared for each protein in 2X binding buffer (50 mM Tris-HCl pH 7.5, 200 mM NaCl, 2 mM MgCl_2_), and 2.5 fmol of labeled RNA diluted in water was denatured at 95°C for 1 min, cooled on ice for 2 min, and combined with an equal volume of 2X binding buffer. The RNA (5 μl) was then combined with the protein serial dilutions for a total reaction volume of 10 μl in 1X binding buffer (resulting in a final RNA concentration of 1.25 nM/reaction and a final protein concentration range of approximately 0-5.5 μM). Reactions were incubated at 37°C for 15 mins and then 5 μl of loading buffer (48% glycerol, 0.01% bromphenol blue) was added to each reaction, and the samples were separated on a native 5% acrylamide gel run at 200 V at 4°C for 3 h with 0.5X TBE buffer (50 mM Tris, 50 mM Boric Acid, 1 mM EDTA pH 8.0). The gels were exposed to a phosphor screen overnight at -20°C and visualized using a BioRad Personal Molecular Imager.

Band intensity was quantified using BioRad Image Lab Software 6. The fraction bound RNA was calculated from the band intensities as bound/(bound+unbound). Fraction bound RNA was then plotted versus concentration of Hdp1_opt_ protein and a hyperbolic equation derived from a one-site binding model was fitted to the data to estimate *K*_d_ values: Fraction bound = *B*_*max*_ [Hdp1_opt_]/(*K*_d_ + [Hdp1_opt_]) + *C*, where *B*_*max*_ represents the maximum fraction bound, *K*_d_ the dissociation constant, and *C* the observed fraction bound at 0 μM Hdp1_opt_. Alternatively, a model accounting for positive cooperativity was also fitted to the data: Fraction bound = *B*_*max*_ [Hdp1_opt_]^*h*^/(*K*_d*h*_ + [Hdp1_opt_]^*h*^) + *C*, where *h* is the Hill coefficient. The latter equation accounts for a model in which two Hdp1_opt_ proteins can bind to one RNA molecule, and where the affinity for the second Hdp1_opt_ is higher than that for the first one. Curve fitting was performed with Prism 9 (GraphPad Software). Refer to Fig. S3 for the fitted parameters for each model. EMSAs were repeated independently three or more times for each protein; representative gels are shown. The quantified values reported represent the mean of three or more independent experimental replicates; the error bars represent the standard error (Fig. S3).

### RNase T1 probing assays

Binding reactions with the radiolabeled full-length *his* operator RNA and the Hdp1_opt_ protein were carried out as described above (using final protein concentrations of 0 and 5.5 μM). Following incubation, 0.05 U of RNase T1 (Ambion) was added, and the reactions were incubated at 37°C for an additional 10 mins. Cleavage was stopped with the addition of 5.5 μl 0.1 M EDTA pH 8.0 (0.33 mM final concentration), and RNA fragments were recovered via ethanol precipitation and resuspended in 10 μl water and 10 μl Gel Loading Buffer II (Ambion). The OH ladder was generated by incubating the RNA in 1X Alkaline Hydrolysis Buffer (Ambion) at 95°C for 12 mins. To generate the denaturing T1 ladder, the RNA was combined with 1X Sequencing Buffer (Ambion), denatured at 95°C for 1 min, cooled on ice, and then incubated with 0.1 U RNase T1 (Ambion) at 37°C for 5 min. Following incubation, ladder reactions were combined with an equal volume of Gel Loading Buffer II (Ambion) and kept on ice. Prior to gel electrophoresis, all samples were denatured at 95°C for 1 min, cooled on ice, and then 5 μl were loaded on an 8% denaturing polyacrylamide gel and run at 30 W at room temperature with 1X TBE (100 mM Tris, 100 mM Boric Acid, 2 mM EDTA pH 8.0). Gels were dried, exposed to phosphor screens for 24-48 h, and visualized using a BioRad Personal Molecular Imager. RNase T1 probing assays were repeated independently three times with similar results; a representative gel is shown.

## ACKNOWLEDGEMENTS

This work was supported by grants from the Wallenberg Foundation (grant 2015.0069 to DIA, grant 2017.0071 to Leif Andersson for the proteomics work) and the Swedish Research Council (grant 2017-01527 to DIA, grant 2019-00666 to MK, grant 2020-04395 to PJ). The funders had no role in study design, data collection and analysis, decision to publish, or preparation of the manuscript. The authors would like to thank Michelle Meyer, Gerhart Wagner, and Omar Warsi for the helpful feedback and suggestions and Cedric Romilly and Thomas Stenum for their assistance with the *in vitro* experiments.

## MATERIALS AVAILABILITY STATEMENT

All newly created materials generated in this study are available upon request.

## AUTHOR CONTRUBUTIONS

AMB, SS, DIA, and MK designed the experiments. AMB, SS, WY, JJH, ML, and MK performed the experiments. AMB, SS, WY, EH, PJ, and MK analyzed data. AMB, DIA, and MK wrote the manuscript with input from all coauthors.

## CONFLICT OF INTEREST

SS is currently employed by CytaCoat (Stockholm, Sweden), WY is currently employed by Sprint Bioscience (Huddinge, Sweden), and DIA consults for Sysmex Europe GmbH (Norderstedt, Germany). These companies were not involved in study design, data collection and analysis, decision to publish, or preparation of the manuscript.

## SUPPLEMENTARY FIGURES

**Supplementary Figure S1:**
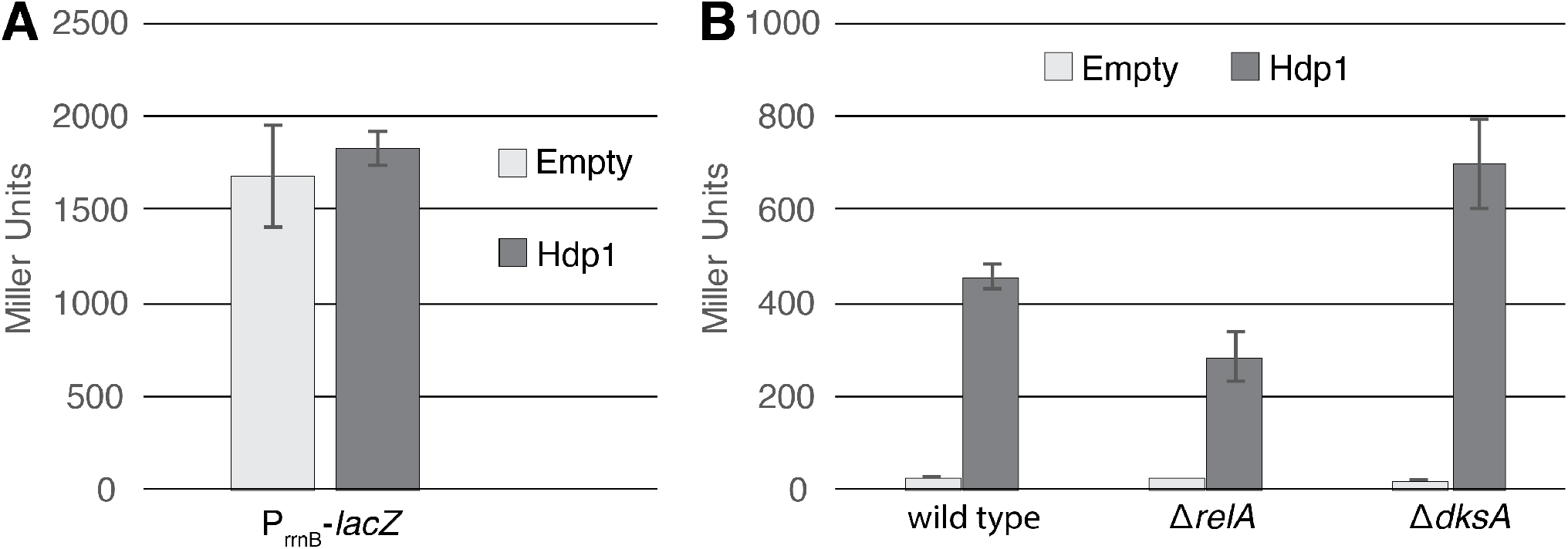
Hdp regulatory activity is independent of the stringent response. (A) β-galactosidase activity (in Miller Units) of the Δ*serB* strain carrying a *lacZ* transcriptional fusion under the control of the *rrnB* P1 promoter and accompanying regulatory sequence upon expression of Hdp1 versus the empty plasmid control. (B) β-galactosidase activity (in Miller Units) of the Δ*serB* strain containing the full-length P_*hisL*_-*his*_operator_-*lacZ* reporter (denoted as “wild type”) and deletions of select stringent response genes upon expression of Hdp1 versus the empty plasmid control. The values reported represent the mean of three or more independent experimental replicates; error bars represent the standard deviation.

**Supplementary Figure S2:**
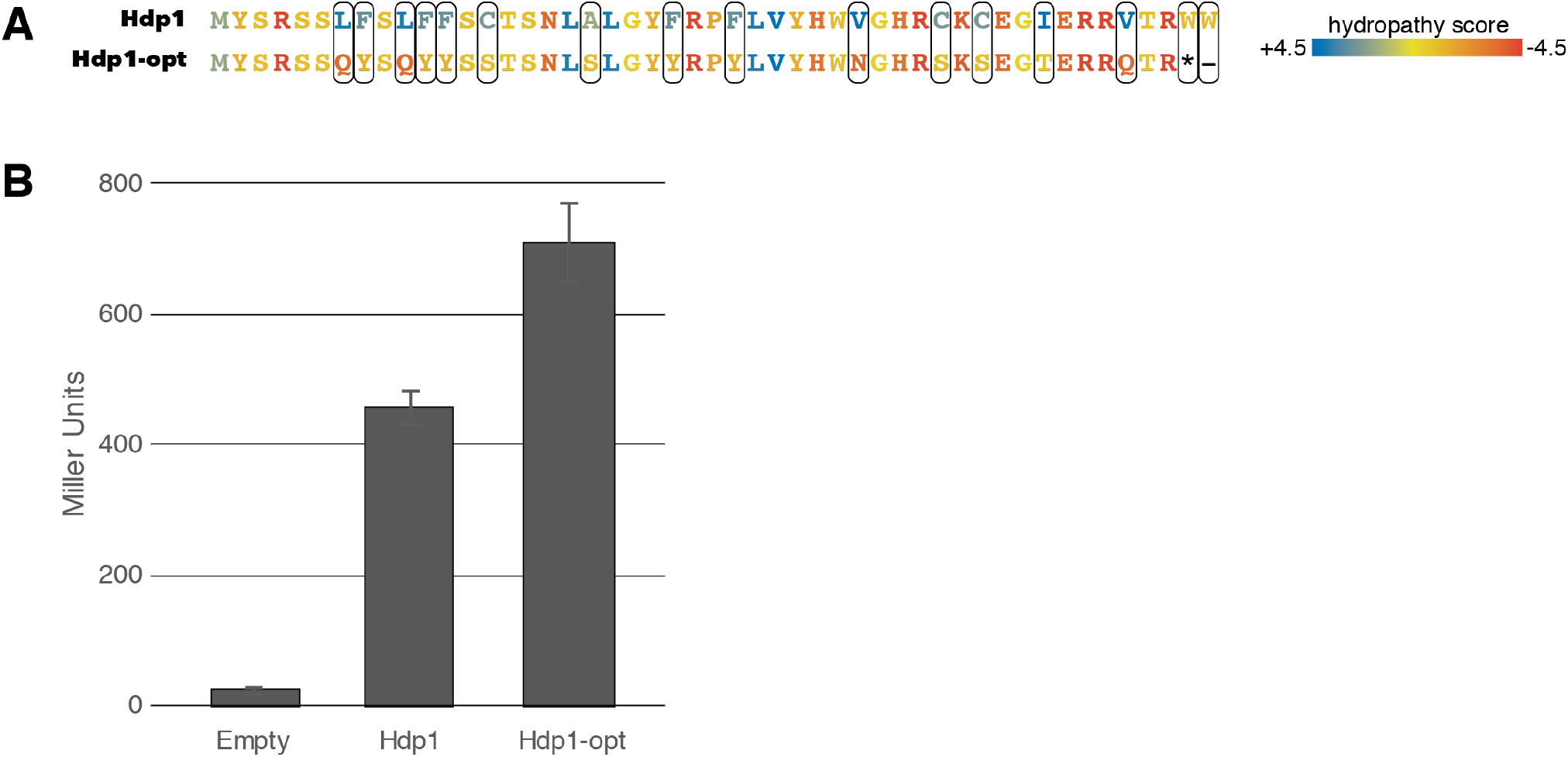
Characterization of Hdp1_opt_, the Hdp1 variant optimized for increased water solubility. (A) Amino acid changes that generated the Hdp1_opt_ variant with improved water solubility. Colors represent the hydropathy score of each amino acid. The grand average of hydropathy (GRAVY) was reduced from -0.043 (Hdp1) to -1.46 (Hdp1_opt_). (B) β-galactosidase activity (in Miller Units) of strains carrying the full-length P_*hisL*_-*his*_operator_-*lacZ* reporter upon expression of the originally isolated Hdp1 protein or the optimized Hdp1_opt_ variant versus the empty plasmid control. The values reported represent the mean of three or more independent experimental replicates; error bars represent the standard deviation.

**Supplementary Figure 3:**
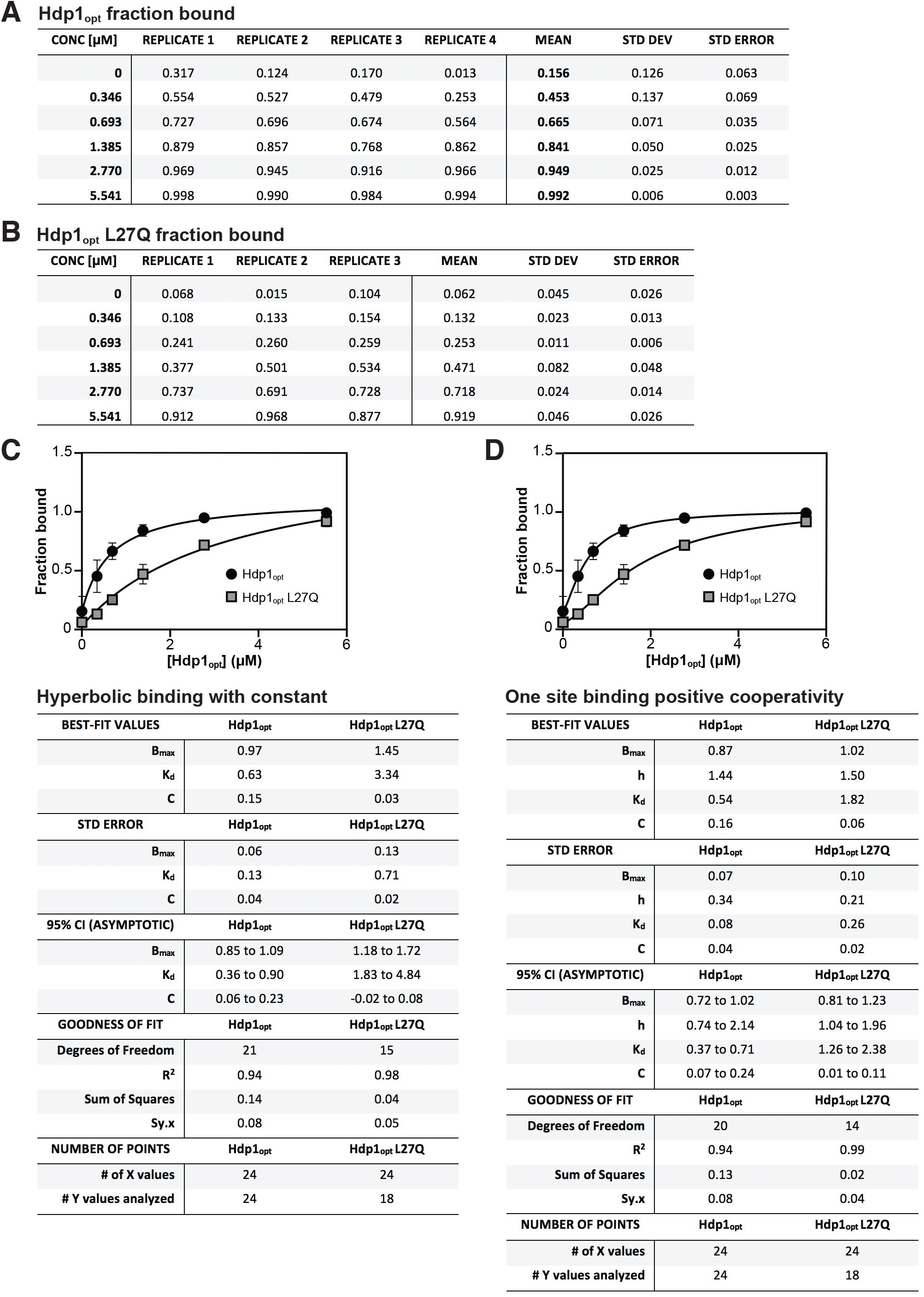
EMSA data and parameters from fitting of different binding models to the EMSA data. Fraction of full-length *his* operator RNA bound to Hdp1_opt_ (A) and Hdp1_opt_ L27Q (B) as quantified and calculated from the EMSA gels. (C) Binding curve and parameters for the EMSA data fitted to a standard 1:1 hyperbolic binding model. (D) Binding curve and parameters for the EMSA data fitted to a model accounting for positive cooperativity. *B*_*max*_ represents the maximum fraction bound, *K*_d_ the dissociation constant, *C* the observed fraction bound at 0 μM Hdp1_opt_, and *h* is the Hill coefficient. Curve fitting was performed with Prism 9 (GraphPad Software); see Materials and Methods for the equations used.

## SUPPLEMENTARY TABLES

**Supplementary Table S1:** KEIO deletion strains screened with the random sequence libraries.

**Supplementary Table S2**: Oligonucleotides used in this study.

**Supplementary Table S3:** *lacZ* reporter strains used in this study; all strains are derived from *Escherichia coli* BW25113.

## FIGURE SUPPLEMENT LEGENDS

**Figure 3A – Figure Supplement 1:** EMSA of the full-length *his* operator RNA in the presence of increasing concentrations of Hdp1_opt_. Uncropped gel from Hdp1_opt_ experimental replicate 1.

**Figure 3A – Figure Supplement 2:** EMSA of the full-length *his* operator RNA in the presence of increasing concentrations of Hdp1_opt_. Uncropped gel from Hdp1_opt_ experimental replicates 2 and 3. Hdp1_opt_ experimental replicate 2 is included in the main text figure as a representative EMSA gel image.

**Figure 3A – Figure Supplement 3:** EMSA of the full-length *his* operator RNA in the presence of increasing concentrations of Hdp1_opt_ (left) or the Hdp1_opt_ L27Q mutant (right). Uncropped gel from Hdp1_opt_ experimental replicate 4 and Hdp1_opt_ L27Q experimental replicate 1. Hdp1_opt_ L27Q experimental replicate 1 is included in the main text figure as a representative EMSA gel image.

**Figure 3A – Figure Supplement 4:** EMSA of the full-length *his* operator RNA in the presence of increasing concentrations of the Hdp1_opt_ L27Q mutant. Uncropped gel from Hdp1_opt_ L27Q experimental replicates 2 and 3.

**Figure 3C – Figure Supplement 1:** Uncropped RNase T1 probing gel for the full-length *his* operator RNA in the absence and presence of Hdp1_opt_ (0, 0.69, and 5.5 μM). NR denotes RNA subject to no reaction, OH indicates partial alkaline hydrolysis, and T1 is an RNase T1 digest of the RNA under denaturing conditions used to map the RNA sequence. (i) Probing reactions incubated with 0.05 U T1 RNase for 5 minutes. (ii) Probing reactions incubated with 0.01 U T1 RNase for 5 minutes. (iii) Probing reactions incubated with 0.05 U T1 RNase for 10 minutes. Numbering of G nucleotides is shown on the left. Arrows highlight changes in RNA cleavage in the presence of Hdp1_opt_: black arrows indicate nucleotides with increased cleavage and white arrows indicate reduced cleavage. Sequence and/or structure characteristics of the *his* operator RNA are also indicated on the right (i.e. the Shine-Dalgarno sequence (SD), start codon (AUG), and stop codon (UAG) of the *hisL* leader peptide coding sequence). The reactions from (iii) are included in the main text figure as a representative T1 RNase probing gel image. All reaction sets were performed with independent RNA and protein dilutions. Contrast has been adjusted from the original gel image so that regions of interest can be labeled.

## Notes

### Competing Interest Statement

The authors have declared no competing interest.

## REFERENCES

Alix, Eric, and Anne Béatrice Blanc-Potard. 2009. “Hydrophobic Peptides: Novel Regulators within Bacterial Membrane.” Molecular Microbiology 72(1):5–11. doi: 10.1111/j.1365-2958.2009.06626.x.

Andrews, Shea J., and Joseph A. Rothnagel. 2014. “Emerging Evidence for Functional Peptides Encoded by Short Open Reading Frames.” Nature Reviews Genetics 15(3):193–204. doi: 10.1038/nrg3520.

Artz, S. W., and J. R. Broach. 1975. “Histidine Regulation in Salmonella Typhimurium: An Activator Attenuator Model of Gene Regulation.” Proc Natl Acad Sci U S A 72(9):3453–57. doi: 10.1073/pnas.72.9.3453.

Aseev, L. V., L. S. Koledinskaya, and I. V. Boni. 2014. “Dissecting the Extended ‘-10’ Escherichia Coli RpsB Promoter Activity and Regulation in Vivo.” Biochemistry (Moscow) 79(8):776–84. doi: 10.1134/S0006297914080057.

Baba, T., T. Ara, M. Hasegawa, Y. Takai, Y. Okumura, M. Baba, K. A. Datsenko, M. Tomita, B. L. Wanner, and H. Mori. 2006. “Construction of Escherichia Coli K-12 in-Frame, Single-Gene Knockout Mutants: The Keio Collection.” Mol Syst Biol 2:2006 0008. doi: 10.1038/msb4100050.

Baek, J., J. Lee, K. Yoon, and H. Lee. 2017. “Identification of Unannotated Small Genes in Salmonella.” G3 (Bethesda) 7(3):983–89. doi: 10.1534/g3.116.036939.

Banerjee, Sharmistha, Jisha Chalissery, Irfan Bandey, and Ranjan Sen. 2006. “Rho-Dependent Transcription Termination: More Questions than Answers.” J Microbiol 44(1):11–22.

Battesti, A., N. Majdalani, and S. Gottesman. 2015. “Stress Sigma Factor RpoS Degradation and Translation Are Sensitive to the State of Central Metabolism.” Proc Natl Acad Sci U S A 112(16):5159–64. doi: 10.1073/pnas.1504639112.

Benjamini, Y., and Y. Hochberg. 1995. “Controlling the False Discovery Rate - a Practical and Powerful Approach to Multiple Testing.” Journal of the Royal Statistical Society Series B-Statistical Methodology 57(1):289–300. doi: DOI 10.1111/j.2517-6161.1995.tb02031.x.

Blasi, F., and C. B. Bruni. 1981. “Regulation of the Histidine Operon: Translation-Controlled Transcription Termination (a Mechanism Common to Several Biosynthetic Operons).” Curr Top Cell Regul 19:1–45. doi: 10.1016/b978-0-12-152819-5.50018-x.

Carvunis, A. R., T. Rolland, I. Wapinski, M. A. Calderwood, M. A. Yildirim, N. Simonis, B. Charloteaux, C. A. Hidalgo, J. Barbette, B. Santhanam, G. A. Brar, J. S. Weissman, A. Regev, N. Thierry-Mieg, M. E. Cusick, and M. Vidal. 2012. “Proto-Genes and de Novo Gene Birth.” Nature 487(7407):370–74. doi: 10.1038/nature11184.

Chojnacki, S., A. Cowley, J. Lee, A. Foix, and R. Lopez. 2017. “Programmatic Access to Bioinformatics Tools from EMBL-EBI Update: 2017.” Nucleic Acids Res 45(W1):W550–53. doi: 10.1093/nar/gkx273.

Cox, J., I. Matic, M. Hilger, N. Nagaraj, M. Selbach, J. V Olsen, and M. Mann. 2009. “A Practical Guide to the MaxQuant Computational Platform for SILAC-Based Quantitative Proteomics.” Nat Protoc 4(5):698–705. doi: 10.1038/nprot.2009.36.

D’Lima, Nadia G., Alexandra Khitun, Aaron D. Rosenbloom, Peijia Yuan, Brandon M. Gassaway, Karl W. Barber, Jesse Rinehart, and Sarah A. Slavoff. 2017. “Comparative Proteomics Enables Identification of Nonannotated Cold Shock Proteins in E. Coli.” Journal of Proteome Research 16(10):3722–31. doi: 10.1021/acs.jproteome.7b00419.

Digianantonio, K. M., and M. H. Hecht. 2016. “A Protein Constructed de Novo Enables Cell Growth by Altering Gene Regulation.” Proc Natl Acad Sci U S A 113(9):2400–2405. doi: 10.1073/pnas.1600566113.

Dinger, M. E., K. C. Pang, T. R. Mercer, and J. S. Mattick. 2008. “Differentiating Protein-Coding and Noncoding RNA: Challenges and Ambiguities.” PLoS Comput Biol 4(11):e1000176. doi: 10.1371/journal.pcbi.1000176.

Dunker, A. Keith, Marc S. Cortese, Pedro Romero, Lilia M. Iakoucheva, and Vladimir N. Uversky. 2005. “Flexible Nets: The Roles of Intrinsic Disorder in Protein Interaction Networks.” FEBS Journal 272(20):5129–48. doi: 10.1111/j.1742-4658.2005.04948.x.

Duval, Mélodie, and Pascale Cossart. 2017. “Small Bacterial and Phagic Proteins: An Updated View on a Rapidly Moving Field.” Current Opinion in Microbiology 39:81–88. doi: 10.1016/j.mib.2017.09.010.

Dyson, H. Jane. 2016. “Making Sense of Intrinsically Disordered Proteins.” Biophysical Journal 110(5):1013–16. doi: 10.1016/j.bpj.2016.01.030.

El-Gebali, S., J. Mistry, A. Bateman, S. R. Eddy, A. Luciani, S. C. Potter, M. Qureshi, L. J. Richardson, G. A. Salazar, A. Smart, E. L. L. Sonnhammer, L. Hirsh, L. Paladin, D. Piovesan, S. C. E. Tosatto, and R. D. Finn. 2019. “The Pfam Protein Families Database in 2019.” Nucleic Acids Res 47(D1):D427–32. doi: 10.1093/nar/gky995.

Fisher, M. A., K. L. McKinley, L. H. Bradley, S. R. Viola, and M. H. Hecht. 2011. “De Novo Designed Proteins from a Library of Artificial Sequences Function in Escherichia Coli and Enable Cell Growth.” PLoS One 6(1):e15364. doi: 10.1371/journal.pone.0015364.

Fu, Yang, Kaila Deiorio-Haggar, Jon Anthony, and Michelle M. Meyer. 2013. “Most RNAs Regulating Ribosomal Protein Biosynthesis in Escherichia Coli Are Narrowly Distributed to Gammaproteobacteria.” Nucleic Acids Research 41(6):3491–3503. doi: 10.1093/nar/gkt055.

Garai, Preeti, and Anne Blanc-Potard. 2020. “Uncovering Small Membrane Proteins in Pathogenic Bacteria: Regulatory Functions and Therapeutic Potential.” Molecular Microbiology 114(5):710–20. doi: 10.1111/mmi.14564.

Gatto, L., and K. S. Lilley. 2012. “MSnbase-an R/Bioconductor Package for Isobaric Tagged Mass Spectrometry Data Visualization, Processing and Quantitation.” Bioinformatics 28(2):288–89. doi: 10.1093/bioinformatics/btr645.

Grisolia, V., A. Riccio, and C. B. Bruni. 1983. “Structure and Function of the Internal Promoter (HisBp) of the Escherichia Coli K-12 Histidine Operon.” J Bacteriol 155(3):1288–96. doi: 10.1128/jb.155.3.1288-1296.1983.

Hemm, M. R., B. J. Paul, T. D. Schneider, G. Storz, and K. E. Rudd. 2008. “Small Membrane Proteins Found by Comparative Genomics and Ribosome Binding Site Models.” Mol Microbiol 70(6):1487–1501. doi: 10.1111/j.1365-2958.2008.06495.x.

Hemm, Matthew R., Jeremy Weaver, and Gisela Storz. 2020. “ Escherichia Coli Small Proteome.” EcoSal Plus 9(1). doi: 10.1128/ecosalplus.esp-0031-2019.

Hoegler, Kenric J., and Michael H. Hecht. 2016. “A de Novo Protein Confers Copper Resistance in Escherichia Coli.” Protein Science 25:1249–59. doi: 10.1002/pro.2871.

Horler, Richard S. P., and Carin K. Vanderpool. 2009. “Homologs of the Small RNA SgrS Are Broadly Distributed in Enteric Bacteria but Have Diverged in Size and Sequence.” Nucleic Acids Research 37(16):5465–76. doi: 10.1093/nar/gkp501.

Johnston, H. M., W. M. Barnes, F. G. Chumley, L. Bossi, and J. R. Roth. 1980. “Model for Regulation of the Histidine Operon of Salmonella.” Proc Natl Acad Sci U S A 77(1):508–12. doi: 10.1073/pnas.77.1.508.

Joyce, A. R., J. L. Reed, A. White, R. Edwards, A. Osterman, T. Baba, H. Mori, S. A. Lesely, B. O. Palsson, and S. Agarwalla. 2006. “Experimental and Computational Assessment of Conditionally Essential Genes in Escherichia Coli.” J Bacteriol 188(23):8259–71. doi: 10.1128/JB.00740-06.

Jumper, John, Richard Evans, Alexander Pritzel, Tim Green, Michael Figurnov, Olaf Ronneberger, Kathryn Tunyasuvunakool, Russ Bates, Augustin Žídek, Anna Potapenko, Alex Bridgland, Clemens Meyer, Simon A. A. Kohl, Andrew J. Ballard, Andrew Cowie, Bernardino Romera-Paredes, Stanislav Nikolov, Rishub Jain, Jonas Adler, Trevor Back, Stig Petersen, David Reiman, Ellen Clancy, Michal Zielinski, Martin Steinegger, Michalina Pacholska, Tamas Berghammer, Sebastian Bodenstein, David Silver, Oriol Vinyals, Andrew W. Senior, Koray Kavukcuoglu, Pushmeet Kohli, and Demis Hassabis. 2021. “Highly Accurate Protein Structure Prediction with AlphaFold.” Nature 596(7873):583–89. doi: 10.1038/s41586-021-03819-2.

Kacar, Betül, Eva Garmendia, Nurcan Tuncbag, Dan I. Andersson, and Diarmaid Hughes. 2017. “Functional Constraints on Replacing an Essential Gene with Its Ancient and Modern Homologs.” MBio 8(4). doi: 10.1128/mBio.01276-17.

Kasai, T. 1974. “Regulation of the Expression of the Histidine Operon in Salmonella Typhimurium.” Nature 249(457):523–27. doi: 10.1038/249523a0.

Knopp, M., J. S. Gudmundsdottir, T. Nilsson, F. Konig, O. Warsi, F. Rajer, P. Adelroth, and D. I. Andersson. 2019. “De Novo Emergence of Peptides That Confer Antibiotic Resistance.” MBio 10(3):e00837–19. doi: 10.1128/mBio.00837-19.

Knopp, Michael, Arianne M. Babina, Jónína S. Gudmundsdóttir, Martin V Douglass, M. Stephen Trent, and Dan I. Andersson. 2021. “A Novel Type of Colistin Resistance Genes Selected from Random Sequence Space.” PLoS Genetics 17(1):1–19. doi: 10.1371/journal.pgen.1009227.

Kolter, R., and C. Yanofsky. 1982. “Attenuation in Amino Acid Biosynthetic Operons.” Annual Review of Genetics 16:113–34. doi: 10.1146/annurev.ge.16.120182.000553.

Kuznetsova, E., M. Proudfoot, C. F. Gonzalez, G. Brown, M. V Omelchenko, I. Borozan, L. Carmel, Y. I. Wolf, H. Mori, A. V Savchenko, C. H. Arrowsmith, E. V Koonin, A. M. Edwards, and A. F. Yakunin. 2006. “Genome-Wide Analysis of Substrate Specificities of the Escherichia Coli Haloacid Dehalogenase-like Phosphatase Family.” J Biol Chem 281(47):36149–61. doi: 10.1074/jbc.M605449200.

Li, Edwin, William C. Wimley, and Kalina Hristova. 2012. “Transmembrane Helix Dimerization: Beyond the Search for Sequence Motifs.” Biochimica et Biophysica Acta - Biomembranes 1818(2):183–93. doi: 10.1016/j.bbamem.2011.08.031.

Mariani, Valerio, Marco Biasini, Alessandro Barbato, and Torsten Schwede. 2013. “LDDT: A Local Superposition-Free Score for Comparing Protein Structures and Models Using Distance Difference Tests.” Bioinformatics 29(21):2722–28. doi: 10.1093/bioinformatics/btt473.

Milligan, John F., Duncan R. Groebe, Gary W. Witherell, and Olke C. Uhlenbeck. 1987. “Oligoribonucleotide Synthesis Using T7 RNA Polymerase and Synthetic DNA Templates.” Nucleic Acids Research 15(21):8783–98. doi: 10.1093/nar/15.21.8783.

Mirdita, Milot, Sergey Ovchinnikov, and Martin Steinegger. 2021. “ColabFold - Making Protein Folding Accessible to All.” BioRxiv 2021.08.15.456425.

Patrick, W. M., E. M. Quandt, D. B. Swartzlander, and I. Matsumura. 2007. “Multicopy Suppression Underpins Metabolic Evolvability.” Mol Biol Evol 24(12):2716–22. doi: 10.1093/molbev/msm204.

Paul, Brian J., Melanie M. Barker, Wilma Ross, David A. Schneider, Cathy Webb, John W. Foster, and Richard L. Gourse. 2004. “DksA: A Critical Component of the Transcription Initiation Machinery That Potentiates the Regulation of RRNA Promoters by PpGpp and the Initiating NTP.” Cell 118(3):311–22. doi: 10.1016/j.cell.2004.07.009.

Paul, Brian J., Melanie B. Berkmen, and Richard L. Gourse. 2005. “DksA Potentiates Direct Activation of Amino Acid Promoters by PpGpp.” Proceedings of the National Academy of Sciences of the United States of America 102(22):7823–28. doi: 10.1073/pnas.0501170102.

Perez-Riverol, Y., A. Csordas, J. Bai, M. Bernal-Llinares, S. Hewapathirana, D. J. Kundu, A. Inuganti, J. Griss, G. Mayer, M. Eisenacher, E. Perez, J. Uszkoreit, J. Pfeuffer, T. Sachsenberg, S. Yilmaz, S. Tiwary, J. Cox, E. Audain, M. Walzer, A. F. Jarnuczak, T. Ternent, A. Brazma, and J. A. Vizcaino. 2019. “The PRIDE Database and Related Tools and Resources in 2019: Improving Support for Quantification Data.” Nucleic Acids Res 47(D1):D442–50. doi: 10.1093/nar/gky1106.

Prijambada, Irfan D., Tetsuya Yomo, Fumihiro Tanaka, Toshihiro Kawama, Keizo Yamamoto, Akihisa Hasegawa, Yasufumi Shima, Seiji Negoro, and Itaru Urabe. 1998. “Solubility of Artificial Proteins with Random Sequences.” Annals of the New York Academy of Sciences 864:131–35. doi: 10.1111/j.1749-6632.1998.tb10295.x.

Ravnikar, P. D., and R. L. Somerville. 1987. “Genetic Characterization of a Highly Efficient Alternate Pathway of Serine Biosynthesis in Escherichia Coli.” Journal of Bacteriology 169(6):2611–17. doi: 10.1128/jb.169.6.2611-2617.1987.

Riggs, D. L., R. D. Mueller, H. S. Kwan, and S. W. Artz. 1986. “Promoter Domain Mediates Guanosine Tetraphosphate Activation of the Histidine Operon.” Proc Natl Acad Sci U S A 83(24):9333–37. doi: 10.1073/pnas.83.24.9333.

Ring, D., Y. Wolman, N. Friedmann, and S. L. Miller. 1972. “Prebiotic Synthesis of Hydrophobic and Protein Amino Acids.” Proceedings of the National Academy of Sciences of the United States of America 69(3):765–68. doi: 10.1073/pnas.69.3.765.

Ritchie, M. E., B. Phipson, D. Wu, Y. Hu, C. W. Law, W. Shi, and G. K. Smyth. 2015. “Limma Powers Differential Expression Analyses for RNA-Sequencing and Microarray Studies.” Nucleic Acids Res 43(7):e47. doi: 10.1093/nar/gkv007.

Ryder, Sean P., Michael I. Recht, and James R. Williamson. 2008. “Quantitative Analysis of Protein-RNA Interactions by Gel Mobility Shift.” Methods in Molecular Biology 488:99–115. doi: 10.1007/978-1-60327-475-3_7.

Samayoa, J., F. H. Yildiz, and K. Karplus. 2011. “Identification of Prokaryotic Small Proteins Using a Comparative Genomic Approach.” Bioinformatics 27(13):1765–71. doi: 10.1093/bioinformatics/btr275.

Sarkar, G., and S. S. Sommer. 1990. “The ‘Megaprimer’ Method of Site-Directed Mutagenesis.” Biotechniques 8(4):404–7.

Sharan, S. K., L. C. Thomason, S. G. Kuznetsov, and D. L. Court. 2009. “Recombineering: A Homologous Recombination-Based Method of Genetic Engineering.” Nat Protoc 4(2):206–23. doi: 10.1038/nprot.2008.227.

Shevchenko, A., M. Wilm, O. Vorm, and M. Mann. 1996. “Mass Spectrometric Sequencing of Proteins Silver-Stained Polyacrylamide Gels.” Anal Chem 68(5):850–58. doi: 10.1021/ac950914h.

Sievers, F., A. Wilm, D. Dineen, T. J. Gibson, K. Karplus, W. Li, R. Lopez, H. McWilliam, M. Remmert, J. Soding, J. D. Thompson, and D. G. Higgins. 2011. “Fast, Scalable Generation of High-Quality Protein Multiple Sequence Alignments Using Clustal Omega.” Mol Syst Biol 7:539. doi: 10.1038/msb.2011.75.

Smith, B. A., A. E. Mularz, and M. H. Hecht. 2015. “Divergent Evolution of a Bifunctional de Novo Protein.” Protein Sci 24(2):246–52. doi: 10.1002/pro.2611.

Steinberg, Ruth, and Hans Georg Koch. 2021. “The Largely Unexplored Biology of Small Proteins in Pro-and Eukaryotes.” FEBS Journal 288(24):7002–24. doi: 10.1111/febs.15845.

Stephens, J. C., S. W. Artz, and B. N. Ames. 1975. “Guanosine 5’-Diphosphate 3’-Diphosphate (PpGpp): Positive Effector for Histidine Operon Transcription and General Signal for Amino-Acid Deficiency.” Proc Natl Acad Sci U S A 72(11):4389–93. doi: 10.1073/pnas.72.11.4389.

Storz, Gisela, Yuri I. Wolf, and Kumaran S. Ramamurthi. 2014. “Small Proteins Can No Longer Be Ignored.” Annual Review of Biochemistry 83:753–77. doi: 10.1146/annurev-biochem-070611-102400.

Su, Mingming, Yunchao Ling, Jun Yu, Jiayan Wu, and Jingfa Xiao. 2013. “Small Proteins: Untapped Area of Potential Biological Importance.” Frontiers in Genetics 4(DEC):1–9. doi: 10.3389/fgene.2013.00286.

Turnbull, Kathryn Jane, Ievgen Dzhygyr, Søren Lindemose, Vasili Hauryliuk, and Mohammad Roghanian. 2019. “Intramolecular Interactions Dominate the Autoregulation of Escherichia Coli Stringent Factor Rela.” Frontiers in Microbiology 10(AUG):1–12. doi: 10.3389/fmicb.2019.01966.

Tyanova, S., T. Temu, and J. Cox. 2016. “The MaxQuant Computational Platform for Mass Spectrometry-Based Shotgun Proteomics.” Nat Protoc 11(12):2301–19. doi: 10.1038/nprot.2016.136.

Vakirlis, Nikolaos, Omer Acar, Brian Hsu, Nelson Castilho Coelho, S. Branden Van Oss, Aaron Wacholder, Kate Medetgul-Ernar, Ray W. Bowman, Cameron P. Hines, John Iannotta, Saurin Bipin Parikh, Aoife McLysaght, Carlos J. Camacho, Allyson F. O’Donnell, Trey Ideker, and Anne Ruxandra Carvunis. 2020. “De Novo Emergence of Adaptive Membrane Proteins from Thymine-Rich Genomic Sequences.” Nature Communications 11(1). doi: 10.1038/s41467-020-14500-z.

VanOrsdel, Caitlin E., John P. Kelly, Brittany N. Burke, Christina D. Lein, Christopher E. Oufiero, Joseph F. Sanchez, Larry E. Wimmers, David J. Hearn, Fatimeh J. Abuikhdair, Kathryn R. Barnhart, Michelle L. Duley, Sarah E. G. Ernst, Briana A. Kenerson, Aubrey J. Serafin, and Matthew R. Hemm. 2018. “Identifying New Small Proteins in Escherichia Coli.” Proteomics 18(10):1700064. doi: 10.1002/pmic.201700064.

Weaver, Jeremy, Fuad Mohammad, Allen R. Buskirk, and Gisela Storz. 2019. “Identifying Small Proteins by Ribosome Profiling with Stalled Initiation Complexes.” MBio 10(2):1–21.

Weidenbach, Katrin, Miriam Gutt, Liam Cassidy, Cynthia Chibani, and Ruth A. Schmitz. 2021. “Small Proteins in Archaea, a Mainly Unexplored World.” Journal of Bacteriology (September). doi: 10.1128/jb.00313-21.

Wilson, B. A., and J. Masel. 2011. “Putatively Noncoding Transcripts Show Extensive Association with Ribosomes.” Genome Biol Evol 3:1245–52. doi: 10.1093/gbe/evr099.

Yip, S. H., and I. Matsumura. 2013. “Substrate Ambiguous Enzymes within the Escherichia Coli Proteome Offer Different Evolutionary Solutions to the Same Problem.” Mol Biol Evol 30(9):2001–12. doi: 10.1093/molbev/mst105.

Yuan, Peijia, Nadia G. D’Lima, and Sarah A. Slavoff. 2018. “Comparative Membrane Proteomics Reveals a Nonannotated E. Coli Heat Shock Protein.” Biochemistry 57(1):56–60. doi: 10.1021/acs.biochem.7b00864.

Zengel, J. M., and L. Lindahl. 1994. “Diverse Mechanisms for Regulating Ribosomal Protein Synthesis in Escherichia Coli.” Prog Nucleic Acid Res Mol Biol 47:331–70.

Zhang, X., and H. Bremer. 1995. “Control of the Escherichia Coli RrnB P1 Promoter Strength by PpGpp.” Journal of Biological Chemistry 270(19):11181–89. doi: 10.1074/jbc.270.19.11181.

Zhang, X., A. H. Smits, G. B. van Tilburg, H. Ovaa, W. Huber, and M. Vermeulen. 2018. “Proteome-Wide Identification of Ubiquitin Interactions Using UbIA-MS.” Nat Protoc 13(3):530–50. doi: 10.1038/nprot.2017.147.

